# The halophilic archaeon *Halogranum roseipondis* sp. nov. is susceptible to a virus carrying an exceptionally high number of viral tRNA genes

**DOI:** 10.1101/2025.11.07.687168

**Authors:** Mirka Lampi, Erika Nordman, Pavlina Gregorova, Janne J. Ravantti, L. Peter Sarin

## Abstract

Archaea constitute a diverse group of organisms, many of which inhabit extreme environments, such as haloarchaea that dominate hypersaline ecosystems, like solar salterns. Although sampling of solar salterns and other hypersaline environments have resulted in numerous haloarchaeal isolates, including 3 classified and 25 unclassified *Halogranum* species, no complete genome has been reported for any member of this genus. Here we present the first comprehensive study of *Halogranum* sp. SS5-1 isolated from a solar saltern in Samut Sakhon, Thailand. *Hgn*. SS5-1 is a pleomorphic, aerobic heterotroph that thrives in high salinity and moderate temperature and is capable of hydrolysing starch. Its genome consists of a 3.6 Mbp chromosome and seven additional plasmids. Based on our phylogenetic analyses, which establish *Hgn*. SS5-1 as a distinct species, we propose that it will be classified as the novel species *Halogranum roseipondis* sp. nov. SS5-1^T^. Additionally, we report that *Hgn. roseipondis* sp.nov. SS5-1^T^ is infected by *Hagravirus capitaneum* (HGTV-1), the only virus known to infect a *Halogranum* host. HGTV-1 exhibits a unique head-tailed morphology and encodes the largest archaeal virus dsDNA genome known to date, including 34 tRNA encoding genes. Codon usage analysis of the viral genome suggests partial alignment with host preferences, yet the abundance of viral tRNA genes hints to broader roles, potentially including translation modulation. This study establishes *Hgn. roseipondis* and HGTV-1 as a novel virus-host system, opening avenues to explore infection dynamics and the roles of virus-encoded tRNA in archaea.

**Importance:** Archaea that thrive in high-salinity environments are key players in geochemical cycles and the primary producers in their habitat. Despite their ecological significance and importance for the development of novel applications in synthetic biology, haloarchaea remain poorly studied. For example, exploration of haloarchaea is required to understand the evolution of cellular complexity and the molecular mechanisms that allow cells to thrive in harsh environmental conditions. In this study, we characterize a novel archaeon, *Halogranum roseipondis* sp. SS5-1^T^, alongside the infection cycle of its associated virus, *Hagravirus capitaneum*. This tailed myovirus carries an extraordinary set of 34 viral tRNA genes, a feature that opens intriguing questions about virus-host interactions and translational control. Our findings lay the groundwork for future investigations into the expression and function of viral tRNAs in an archaeal model system, thereby enabling the study of archaeal translation and virus-driven modulation of host cellular processes.

## Introduction

Archaea constitute one of the three domains of life and includes a diverse group of extremophiles that thrive in seemingly uninhabitable environments, such as acidic hot springs (1), hydrothermal vents (2), and soda lakes (3). Amongst these extremophiles, haloarchaea inhabit hypersaline environments, in which they are often the dominant microorganisms outcompeting halophilic bacteria and eukaryotes (4). Such hypersaline conditions are encountered in e.g. marine solar saltern ponds, which are man-made environments where seawater is evaporated to crystallize salt. The salinity of these shallow evaporation ponds may vary between that of typical sea water to crystallized salt. Unlike most extreme habitats, evaporation ponds are easily accessible. Hence, they have been extensively sampled and studied, uncovering archaea mainly belonging to the order *Halobacteriales* and families *Haloferacaceae* and *Haloarculaceae* (5–9). *Haloferacaceae* consists of 26 genera, of which *Haloferax* and *Halorubrum* contain 23 and 50 classified species, respectively, whereas most other genera are small and include 3 or fewer species (10, 11). Of these, genus *Halogranum* contains 3 characterized and 26 uncharacterized species; the latter mainly discovered following a systematic search for halophilic viruses from a solar saltern in Samut Sakhon, Thailand (5, 6). Interestingly, many haloarchaeal virus isolates from high salinity environments are reported to have a wide host range; especially species belonging to *Halorubrum* and *Haloarcula* are susceptible to infection by several different virus isolates. However, of the 85 haloarchaeal viruses tested on *Halogranum* isolates so far, only one is known to infect a *Halogranum* host (5, 6).

To date, genus *Halogranum* contains three characterized species, all isolated from solar salterns. The genus was originally created based on *Hgn. rubrum* (12) and later expanded with *Hgn. salarium* (13), *Hgn. amylolyticum* and *Hgn. gelatinilyticum* (14), of which *Hgn. salarium* has recently been phylogenetically reclassified as a strain of *Hgn. rubrum* (15). All characterized *Halogranum* species are extremely halophilic, thriving at a total salinity range of 2.6 to 3.9 M (12–14). In this study, we present the first detailed characterization of the haloarchaeon *Halogranum* sp. SS5-1 (5). We show that *Hgn*. sp. SS5-1 cells are pleomorphic, non-motile, aerobic heterotrophs with optimal growth at 42 °C, pH 6-7.5 in the presence of 3.4 M salt. *Hgn*. sp. SS5-1 utilizes a variety of single carbon sources, and it hydrolyzes starch but not gelatin. Furthermore, we also present the complete genome of *Hgn*. sp. SS5-1—the first complete genome in the *Halogranum* genus—consisting of a 3 617 kbp dsDNA main chromosome and 7 additional plasmids that range in size from 40 kbp to 561 kbp. Using Roary pangenomic analysis, we group *Hgn*. sp. SS5-1 to the *Halogranum* family next to *Haloprofundus* genus. Similarity analysis of the three 16S ribosomal RNA genes and the average nucleotide identity (ANI) analysis further suggests that *Hgn*. sp. SS5-1 constitutes a distinct novel species that is closely related to *Hgn. rubrum* and *Hgn. amylolyticum*. Consequently, we propose that *Hgn*. sp. SS5-1 should be classified as a novel species within the *Halogranum* genus, bearing the designation *Halogranum roseipondis nov*. SS5-1^T^, wherein ‘*rosei*’ refers to the pink color of the species in liquid culture and ‘*pondis*’ to the isolation pond, i.e. solar saltern.

In addition, here we also introduce the first infection cycle characterization for a virus infecting an archaeon from genus *Halogranum*: *Hagravirus capitaneum* (HGTV-1) (5, 16). *Hagravirus capitaneum* belongs to the order *Thumleimavirales* representing the only known species in family *Halomagnusviridae* and *Hagravirus* genus. As for all head-tailed viruses known to date, HGTV-1 has a lytic infection cycle with progeny virions being released to the cell’s surroundings. We show that the HGTV-1 infection cycle takes approximately 10 hours, terminating with a modest amount of progeny virus production. We also demonstrate that HGTV-1 has an unusual morphotype with an icosahedral capsid, connected to the tail by a long and thin neck structure. Interestingly, HGTV-1 has the largest double stranded DNA (dsDNA) genome of any known archaeal virus to date. Reanalysis of tRNA genes established that it carries 34 virus-encoded tRNA (vtRNA) genes covering all amino acids in the universal genetic code, except for histidine. Previous studies have suggested that vtRNAs may bridge biases caused by codon usage differences in the host and virus genomes (17). Our codon usage comparison revealed slight biases that might explain the presence of some of the vtRNA genes in the HGTV-1 genome, but not all of them. Hence, we hypothesize that in addition to bridging codon usage differences, vtRNAs might partake in other infection-supporting roles, such as acting as translational decoys (18), for which additional studies are needed. Importantly, this constitutes the first detailed host-virus characterization of a *Halogranum* infection model, paving the way for further studies on viral tRNA functions in an archaeal host organism.

## Results

### Protologue

*Halogranum* species, *Halogranum roseipondis* SS5-1^T^ sp. nov. *Roseipondis* (ro.se.i.pon′dis. L. adj. roseus, rosy; L. n. pondus, pond; N.L. gen. n. roseipondis, of a rosy pond), referring to the pink coloration of the liquid culture and the hypersaline pond from which this species was isolated.

*Halogranum roseipondis* SS5-1^T^ is a pleomorphic haloarchaeon that grows at near neutral conditions (pH 6.5- 7) in a wide range of salinities (2.7-4.2 M) and temperatures (25-50 °C). *Hgn. roseipondis* SS5-1^T^ has an aerobic metabolism, it utilizes a variety of carbon sources including fructose, glucose, glycerol, sucrose, Na-succinate, Na-lactate, and Na-acetate, and it hydrolyzes starch but not gelatin. It is positive for catalase and oxidase activity (Table 1), and is sensitive to rifampicin, mevinolin and novobiosin (Table 1). *Hgn. roseipondis* SS5-1^T^ produces a red/pink pigment and may feature extracellular surface appendages that seemingly connect the cells (Fig. 1). *Hgn. roseipondis* SS5-1^T^ can be infected with *Hagravirus capitaneum* HGTV-1.

**Table 1.**
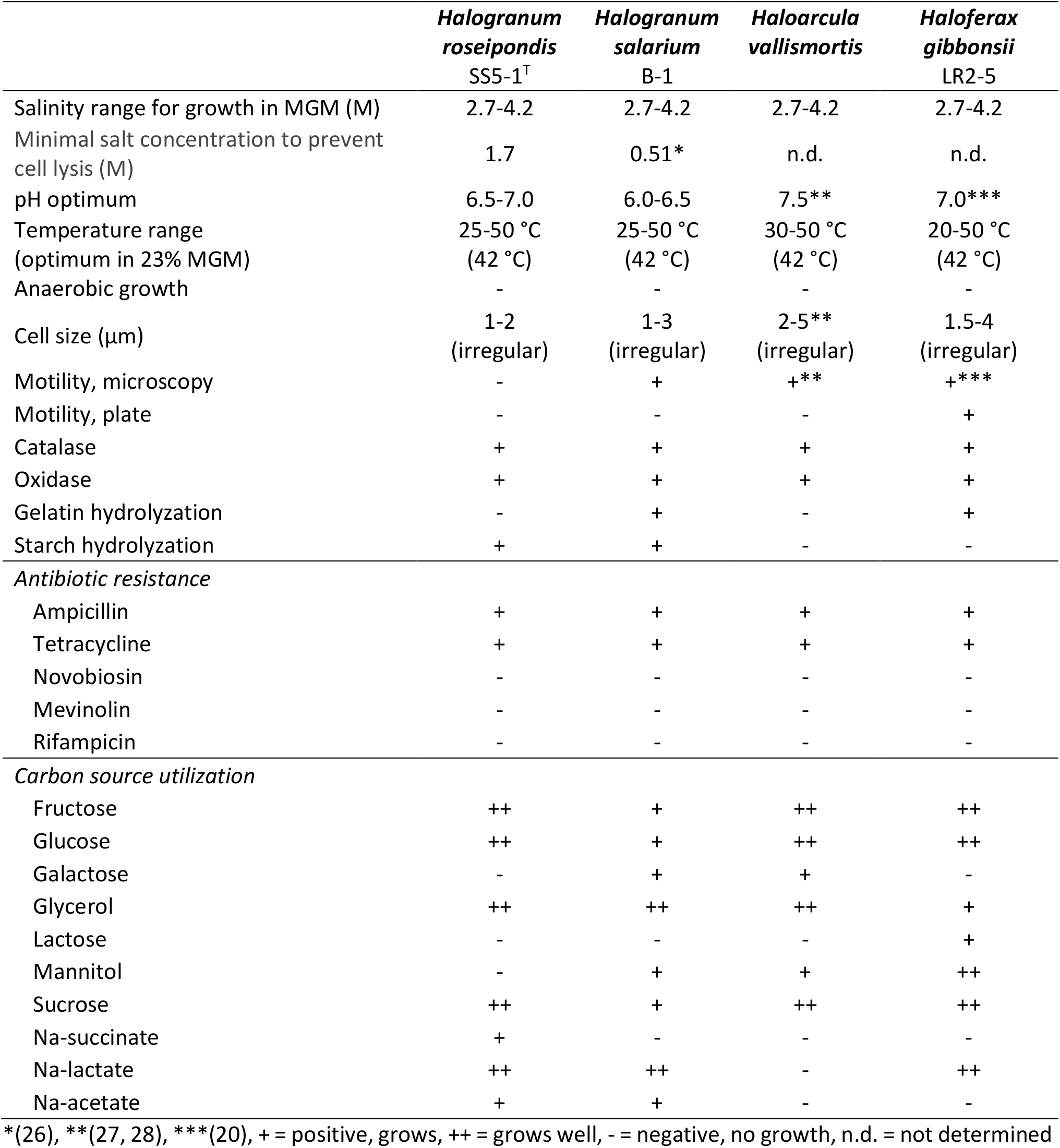
Characteristics of *Hgn. roseipondis* SS5-1^T^ and reference strains.

**Figure 1.**
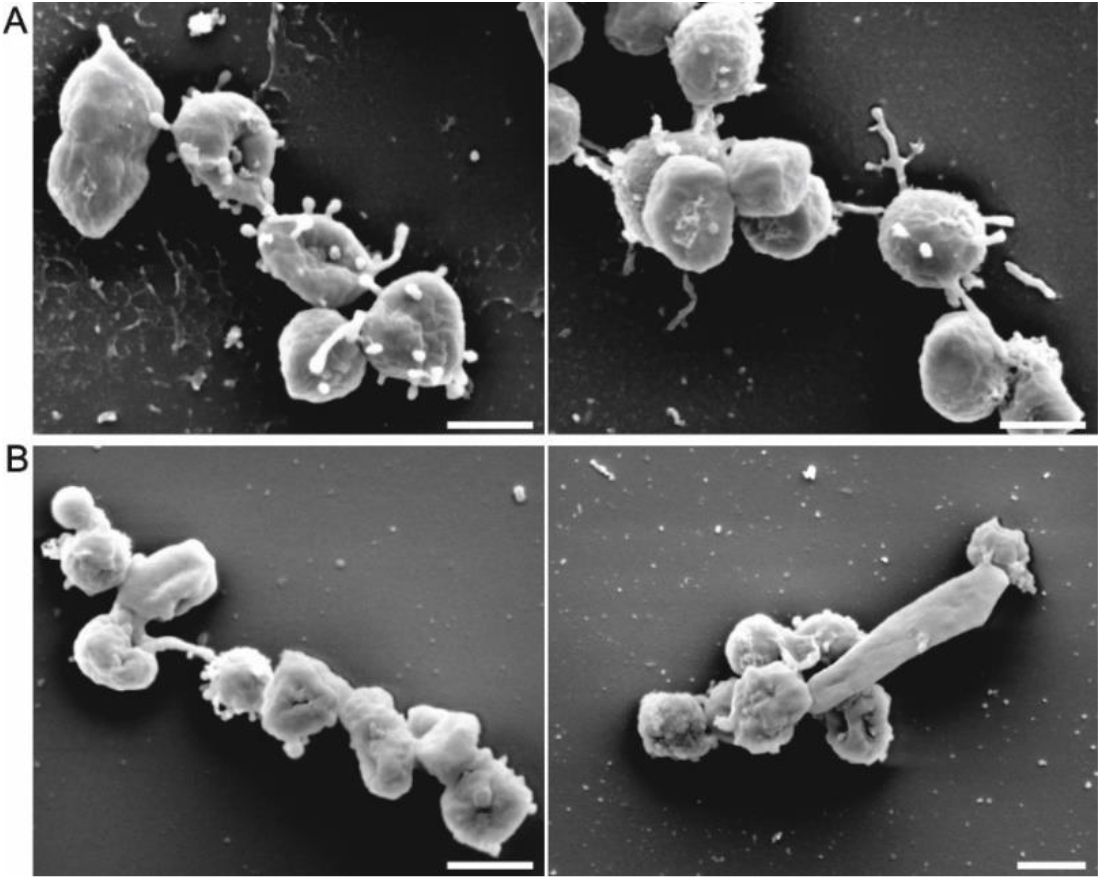
*Halogranum roseipondis* SS5-1^T^cells are irregularly shaped. A) Scanning electron micrographs of *Hgn. roseipondis* grown to OD_600_ of 0.6. B) *Hgn. roseipondis* cells grown to OD_600_ of 1. The size bar in all panels correspond to 1 µm.

*Halogranum roseipondis* SS5-1^T^ belongs to the genus *Halogranum* within the family *Halobacteriaceae*, class *Halobacteria*, domain *Archaea*. Phylogenetic analysis based on 16S rRNA gene sequences places *Hgn. roseipondis* next to *Halogranum amylolyticum* and *Halogranum rubrum*. OrthoANI analysis reveals the ANI-value to the closest relative, *Halogranum amylolyticum*, to be 92.28. The *Hgn. roseipondis* SS5-1^T^ genome consists of a 3 617 299 bp main chromosome (GC% 65.51) and seven additional plasmids ranging from 40 297 to 560 995 bp in size (Fig. 2).

**Figure 2.**
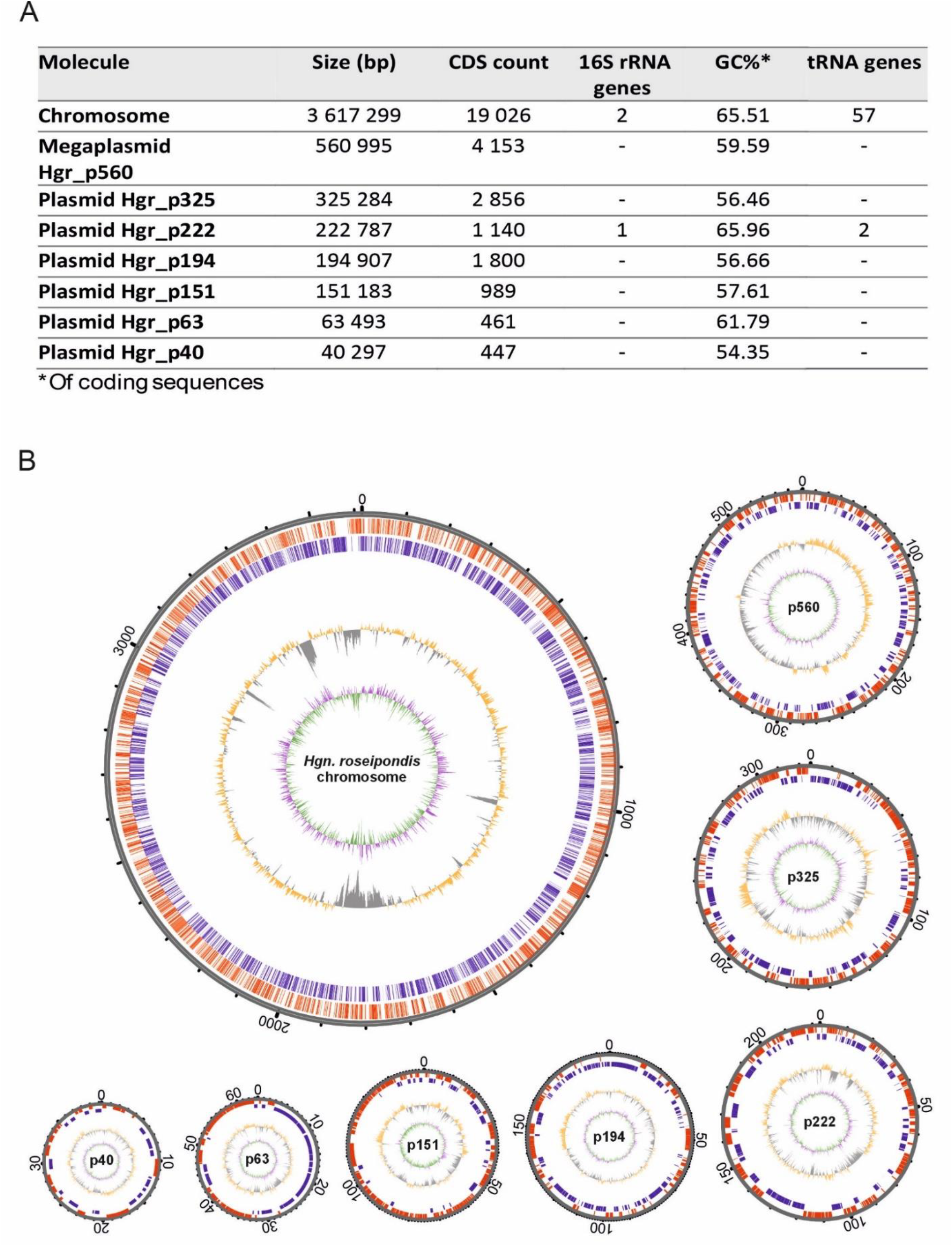
The *Halogranum roseipondis* SS5-1^T^ genome consists of a large chromosome and seven additional DNA molecules. A) Key numerical details of the eight *Hgn. roseipondis* genome molecules. B) Circular genome maps of each genome molecule. From outside to center: coding DNA sequence (CDS) forward strand and CDS reverse strand presented in red and blue, respectively. The middle circle (yellow/grey) represents GC content distribution, and the innermost circle (green/purple) shows the GC skew. Size annotations on the outer ring of the map are in kb. To facilitate legibility, circular map sizes are not to scale.

### *Halogranum roseipondis* is an extreme halophile able to utilize several single carbon sources

*Halogranum roseipondis* SS5-1^T^ is a non-motile, obligate aerobe that was originally isolated from a salt crystallization pond in Samut Sakhon, Thailand (Table 1, Fig. S1) (5). *Hgn. roseipondis* SS5-1^T^ (hereafter referred to as *Hgn. roseipondis*) forms round (∼2 mm in diameter) and shiny colonies with a distinct pink coloration, possibly due to carotenoid production—a common trait in many haloarchaea (19). However, pigment production diminishes as growth temperatures exceed 45 °C, with colonies acquiring a yellowish/light orange color (Fig. S2). We determined that *Hgn. roseipondis* grows at a temperature range from 25 °C to 45 °C with an optimum at 42 °C when assayed in liquid microcultures (Table 1, Fig. S3). However, growth can be sustained at 50 °C both on solid media and in liquid flask cultures, although aggregates frequently appear at >45 °C, suggesting that the cells are at their survivability limit. Next, we characterized the impact of salinity at different temperatures. *Hgn. roseipondis* grows optimally in 23% MGM (3.4 M total salts) at 42 °C (Table 1, Fig. S3), whereas a ∼2 M (12% MGM) minimal salt concentration is required for growth. *Hgn. roseipondis* cells remain intact in 10% (w/v) NaCl (1.71 M) solutions, but lysis is immediate in ddH_2_O. Conversely, growth was still detected at 28% MGM (4.1 M total salts), although liquid cultures are difficult to handle at such high salinity due to salt crystallization. Interestingly, we also observed that 12-18% MGM (1.8- 2.6 M total salinity) concentrations were more favorable at lower (<37 °C) growth temperatures (Fig. S3). We also established that *Hgn. roseipondis* has a narrow pH range of 6.0-7.5 with a preference for neutral towards slightly acidic environments (Table 1, Fig. S4). Furthermore, *Hgn. roseipondis* can grow on several single carbon sources and it hydrolyzes starch but not gelatin (Table 1, Fig. S5). We also concluded that growth is inhibited by novobiocin, mevinolin and rifampicin, whereas no inhibition was observed with other antibiotics tested (Table 1, Fig. S6). Hence, we conclude that optimum laboratory growth conditions for *Hgn. roseipondis* are achieved on rich 23% MGM media at 42 °C and pH 6.5-7.

### *Hgn. roseipondis* cells are irregularly shaped and feature club-like surface structures

Our initial transmission electron microscopy (TEM) findings suggested that *Hgn. roseipondis* might have an irregular cell morphology (Fig. S7). Similar observations have been made for other haloarchaea, such as *Haloferax gibbonsii*, where cell morphology changes are tightly linked to the growth phase (20). Consequently, we set out to investigate if such a connection is present in *Hgn. roseipondis* by examining its cellular morphology at two growth phases: early exponential growth phase (OD_600_ 0.6) cells are irregularly shaped, although we noticed a slight prevalence towards coccoidal cells with a diameter of ∼1 µm (Fig. 1A), whereas cells at a later exponential growth phase (OD_600_ 1) appeared more irregular, elongated, and angular with varying sizes being observed (Fig. 1B). Irrespective of growth phase, some cells feature club-like structures/surface protrusions, which seem to connect some of the cells together (Fig. 1). It has previously been reported that archaea express unique attachment and anchoring structures, like hami (21), cannulae (22, 23), and pili, which facilitate adhesion and biofilm formation (24), as well as archaeal flagella, *i*.*e*. archaella, for motility (25). However, the surface structures of *Hgn. roseipondis* do not directly resemble any of these previously reported structures and their function remains to be determined.

### Phylogenetic analyses suggest that *Hgn. roseipondis* constitutes a novel *Halogranum* species

The *Hgn. roseipondis* genome was sequenced using a PacBio Sequel II sequencer and assembled using Pacbio’s Microbial Genome Analysis application. A total of 10 761 PacBio HiFi sequence reads was obtained with a N50 read length being 16 828 bp (381-43 667 bp), resulting in an average coverage depth of 18.7- 44.2× (Table S1). *Halogranum roseipondis* genome consists of a large 3 617 kbp chromosome, a 560 kbp megaplasmid, and six additional plasmids ranging between 40 – 325 kbp in size (Fig. 2). The chromosomal GC content is 65.51% and plasmid GC contents differ to some extent, varying between 54.35% and 65.96%.

Next, we determined how the *Hgn. roseipondis* genome relates to other haloarchaea. Roary pangenomic analysis places *Hgn. roseipondis* in the same cluster as other *Halogranum* species when compared to representative type species from all genera within the family *Haloferacaceae*, except for *Candidatus Haloextosymbiotes* (Fig. 3A, Table S2, Fig. S8). This is supported by phylogeny based on 16S rRNA sequence similarity, which places *Hgn. roseipondis* next to *Hgn. rubrum* and *Hgn. amylolyticum*. (Fig. 3B). Notably, two 16S rRNA genes are present in the *Hgn. roseipondis* chromosome and one in the plasmid Hgr_p222 (Fig. 2A). There is a minor intragenomic variance between the two chromosomal 16S rRNA genes (99.7% sequence similarity). Likewise, 99.7% and 99.5% sequence similarities are observed when comparing the plasmid-based 16S rRNA gene to the respective chromosomal genes. Furthermore, average nucleotide identity (ANI) analysis suggests that *Hgn. roseipondis* should be classified as a novel *Halogranum* species as the highest ANI score obtained is 92.28 (to *Hgn. amylolyticum*), which exceeds the 5% threshold and is sufficiently different for *Hgn. roseipondis* to be considered as a novel species (Fig. 3C, Fig. S9) (29, 30).

**Figure 3.**
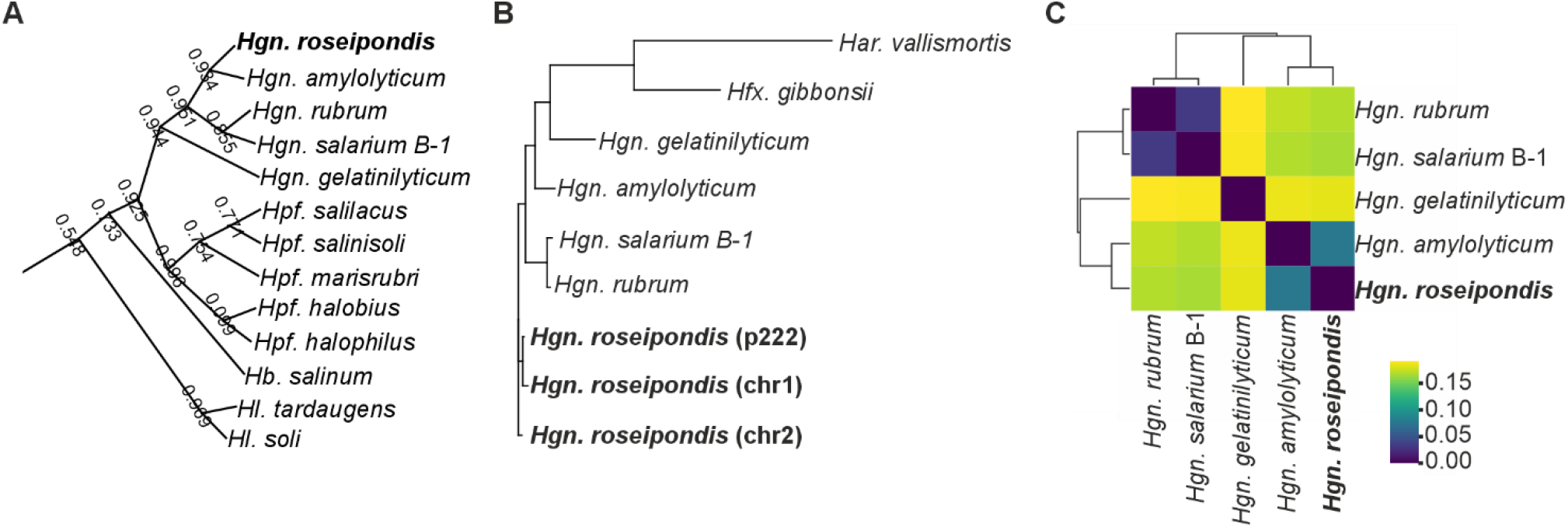
Phylogenetic relationship of *Hgn. roseipondis* SS5-1^T^. A) Roary pangenomic analysis places *Hgn. roseipondis* on the same branch as *Hgn. amylolyticum*. Genera abbreviations: *Haloprofundus* = *Hpf*., *Halobium* = *Hb*. and *Halegenticoccus* = *Hl*. For the complete tree including 125 *Haloferacaceae* species, see Fig. S8. B) Phylogenetic tree of 16S rRNA gene similarity within the genus *Halogranum* and the reference strains used in this study. The location of *Hgn. roseipondis* 16S rRNA genes are presented in brackets; p222= in plasmid Hgr_p222, Chr1= in chromosome, location 1 345 500 -1 346 971, Chr2 = in chromosome, location 2 929 224-2 930 695. C) orthoANI analysis of *Halogranum* species. The color legend represents orthoANI value fractions, where a higher value indicates less similarity. For the complete ANI analysis with 125 *Haloferacaceae* species, see Fig. S9.

### *Hgn. roseipondis* is infected by a virus with an unusual head-tailed morphotype

Many archaeal viruses are known for their extraordinary virion morphologies (31). The HGVT-1 virion has a total length of ∼230 nm and represents a rather ‘classical’ head-tailed morphotype with an icosahedral capsid (∼75 nm in diameter) and a long tail (∼155 nm) (Fig. 4A, B). Nevertheless, the tail structure is exceptional as it is composed of a thin neck and a much wider clamp-like expansion (Fig. 4A, B). Moreover, based on frequent electron microscopy observations, the tail structure seems to easily detach from the virion (Fig. S10), although it is not clear whether this is solely an artefact of sample preparation. We also identified tail assemblies of various lengths, some 3-4× the length of the mature tail structure, which might be non-processed assembly intermediates (Fig. S10). However, neither virion assembly nor the function of the clamp-like expansion are within the scope of this work, and further studies are needed to address these aspects.

**Figure 4.**
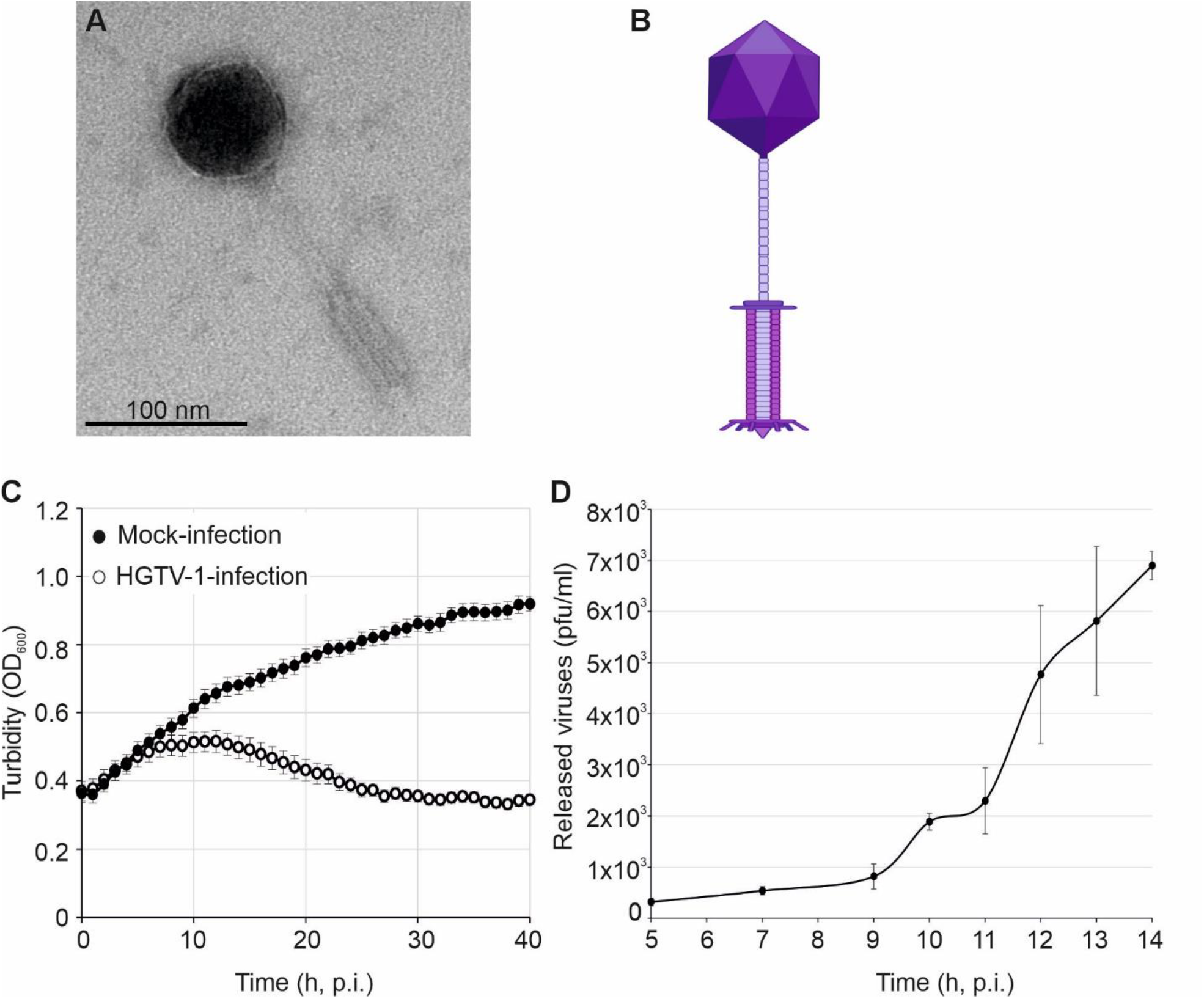
HGTV-1 has an unusual head-tail morphotype and lytic infection cycle. A) A transmission electron micrograph of the HGTV-1 virion shows a thin long neck that connects the tail to the capsid. B) Schematic interpretation of the HGTV-1 virion (based on A). C) Representative growth curves of HGTV-1 infected and mock-infected *Hgn. roseipondis* cultures. Note that the turbidity of the infected culture begins to decrease at 12 h p.i. D) One-step growth curve of HGTV-1 infected *Hgn. roseipondis* cells. Release of progeny virions is observed at 10 h p.i., followed by increasing numbers of released virions. Error bars denote standard deviation (n=3).

HGTV-1 has been previously reported to infect *Har. vallismortis*, albeit with a very low infectivity (5). Our cross-infection studies confirmed these initial results. We also did not observe any infectivity with the reference strains *Hgn. salarium* or *Hfx. gibbonsii* (Fig. S11).

HGTV-1 infects *Hgn. roseipondis* most efficiently at 37 °C using 23% MGM growth medium (Fig. 4). At higher temperatures and/or salinities, the host becomes more resistant to infection, possibly since the conditions are less favorable to virus adsorption, allowing the host culture to outgrow the infection (Fig. S12). The HGTV- 1 infection cycle is lytic and infected cell bursts ∼12 h post infection (p.i.), as is noticeable from the declining turbidity of the host culture (Fig. 4C). However, the infection cycle is not fully synchronized, and nascent virions can be detected as early as 10 h p.i., followed by a subsequent increase in the number of released virions. Based on one-step growth curves, we estimate the burst size after 14h p.i. to be modest, releasing on average only 20 virions per infected cell (Fig. 4D).

### HGTV-1 encodes numerous viral tRNAs of unknown function

HGTV-1 has previously been reported to have a genome size of 143 855 bp (16)—the largest genome of any archaeal virus known to date. In addition, HGTV-1 also carries 36 virus-encoded tRNA (vtRNA) genes including 2 pseudogenes, constituting the highest number of vtRNA genes encoded by an archaeal virus (16). Following a reanalysis of the published genome with tRNAScan-SE 2.0 (32), we identified 34 tRNA genes—3 putative pseudogenes, 2 tRNA genes of undetermined type, and 29 regular tRNA genes with an anticodon that matches the body of the isoacceptor (Table S3). Moreover, one tRNA gene also carries an intron. We also noted that the average GC content of HGTV-1 vtRNAs is somewhat lower (55.2%; range 38.6-63.9%) than that of the host tRNAs (61%; range 50.7-68.2%) (Table S3).

The presence of vtRNA genes is often considered a key adaptation mechanism for bridging potential discrepancies in codon usage and availability between the virus and the host (33). To investigate this further, we performed a tRNA and codon usage analysis for all coding sequences in the *Hgn. roseipondis* and HGTV- 1 genomes, respectively. As expected, the host encodes for tRNAs that match the preferred codons, such as GAC for aspartic acid, TTC for phenylalanine, AAC for asparagine, and TAC for tyrosine. When codon usage is non-preferential between the possible codon options, as is the case for glycine, arginine, and serine, *Hgn. roseipondis* also encodes tRNAs that match the optional codons (Table S3, S4). However, our codon usage comparison between the host and the virus revealed differences in the preferred codons used for glutamic acid, glycine, isoleucine, and glutamine—*Hgn. roseipondis* favors pyrimidine (C/G) as the third codon base whereas purine (A/T) is prevalent at the same position in HGTV-1 (Fig. 5, Table S4). Importantly, these limitations can be overcome for glutamic acid, glycine, and glutamine as HGTV-1 encodes vtRNAs with both pyrimidine and purine at the third base, thus supplementing the translation machinery with tRNAs that match transcripts originating from both genomes. Conversely, this is not the case for isoleucine, where the vtRNA matches the preferred host codon but not that of the virus (Fig. 5A). For asparagine, codon AAC is clearly more dominant in the host whereas codons AAC and AAT are used almost equally in the virus genome. Bearing this in mind, it is surprising that HGTV-1 encodes a vtRNA for ACC only but none for AAT (Fig. 5B). In the case of leucine, HGTV-1 encodes vtRNAs for codons CTC, CTA, TTA, and TTG. CTC is the most prevalent codon in both the host and the virus, whereas optional codons CTA, TTA, and TTG are used more pronouncedly by the virus, suggesting that expression of these vtRNAs would further HGTV-1 protein translation (Fig. 5B). A similar situation is observed for proline and alanine, where the preferred codon is the same for *Hgn. roseipondis* and HGTV-1, but utilization of the A ending codon with matching vtRNA is more prominent in the virus for optional codons (Fig. 5B). Whether viral tRNAs are recognized by the ribosome— given that their sequences differ to some extent from host tRNAs (Fig. S13)—remains an open question for future studies to address.

**Figure 5.**
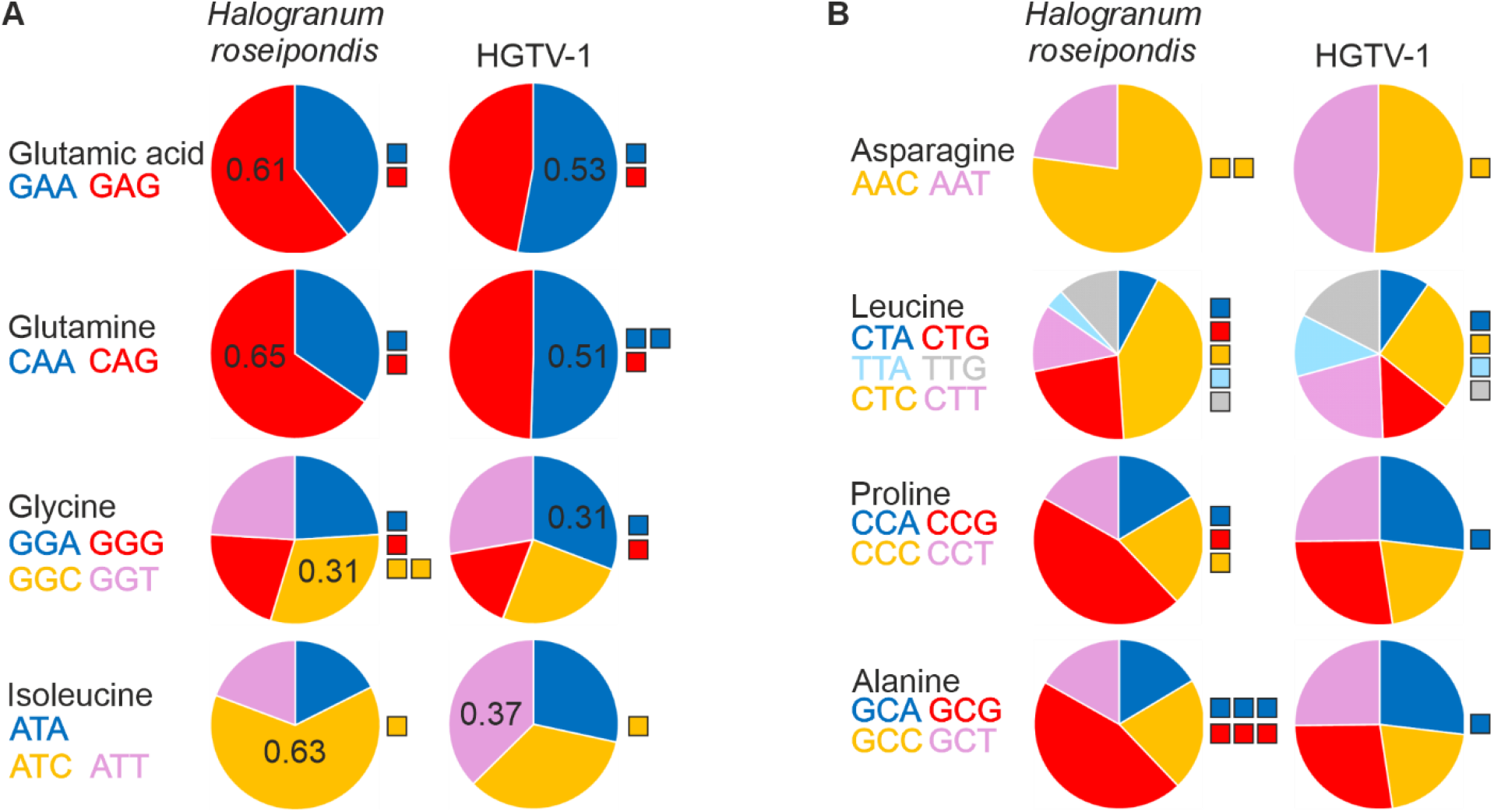
Codon usage analysis of *Hgn. roseipondis* SS5-1^T^ and HGTV-1 coding regions. A) Codon usage comparison in cases where the preferred codon was different in *Hgn. roseipondis* and HGTV-1. The number in each pie chart indicates the fraction of the preferred codon. B) Codon usage comparison of selected amino acids where the preferred codon is the same, but optional codons differ. Pie chart colors correspond to the color of the codons at the left side of each chart. For A) and B), the colored squares next to the charts represent codon matching tRNA genes carried by *Hgn. roseipondis* or HGTV-1, respectively. Each square depicts one tRNA coding gene.

## Discussion

In this study, we provide the first detailed characterization of the physiological and genomic features of *Halogranum* sp. SS5-1 (5), based upon which, we propose that it be classified as the novel species *Halogranum roseipondis* sp. nov. SS5-1^T^. Previously characterized *Halogranum* species have all been isolated from solar salterns and they are known to grow in 2.6-3.9 M NaCl and near-neutral pH conditions (12, 26). The novel species *Hgn. roseipondis* sp. nov. SS5-1^T^ shares these fundamental growth requirements (Table 1) that align with the environmental conditions in solar salterns, where salinity and temperature may fluctuate significantly. Moreover, many haloarchaeal species produce carotenoid pigments that range in color from yellow to red (19). These carotenoids act as photoprotectors and antioxidants, protecting the cell from UV radiation damage and oxidative stress, in addition to which they have also been associated with DNA repair and membrane fluidity control (19). *Hgn. roseipondis* produces a pink/reddish pigment at optimal growth conditions (Fig. S2). However, we did observe loss of pigment production at growth temperatures exceeding 45 °C (Fig. S2), which may be potentially attributed to transcriptional reprogramming, e.g. induction of heat shock responses etc., as the cells reach the limit of their survival.

Many archaea are known to feature unusual cell morphologies (34) as well as undergo growth phase-dependent morphological changes (20, 35). Our microscopic analyses revealed that *Hgn. roseipondis* cells are pleomorphic with a predominantly coccoid morphology at the early exponential growth phase, whereas cell sizes and morphologies diversify when the culture is grown to a later exponential growth phase (Fig. 1). Furthermore, we also observed that *Hgn. roseipondis* cells are occasionally decorated with extracellular appendices, which seemingly link individual cells together (Fig. 1). These structures do not resemble any previously reported archaea-specific extracellular structures like cannulae or hami (21–23), nor have such structures been reported for any other *Halogranum* species. Although elucidating the function of these surface structures falls outside the scope of this study, it is conceivable that they may constitute a previously undescribed component that contributes to e.g. archaeal biofilm formation.

The genome of *Halogranum roseipondis* represents the first complete and fully assembled *Halogranum* genome (Fig. 2). It consists of a large main chromosome, a megaplasmid, and six smaller plasmids (Fig. 2). Plasmids are common in archaea, where they often carry genes that are critical for adaptation and survival. Haloarchaea typically harbour 2-7 plasmids, in addition to which they may have unique extrachromosomal elements, such as megaplasmids or minichromosomes, that are not found in other archaeal species (36). Notably, the 561 kbp megaplasmid of *Hgn. roseipondis* genome (Fig. 2) has a significantly lower GC content than the main chromosome – a feature that is typical for haloarchaea and likewise indicative of the megaplasmid deriving from a different origin (36). Conversely, plasmid Hgr_p222 carries both rRNA and tRNA genes and has a comparable GC-content to the main chromosome. This suggests that despite its relatively small size, Hgr_p222 could be seen as a mini-chromosome, similar to a bacterial chromid, rather than a plasmid (37, 38). The plasmids of *Halogranum roseipondis* may point to a mosaic evolutionary history, potentially enabling integration of features such as extracellular appendices, that are of uncertain origins and have not been documented in haloarchaeal species studied to date. Our phylogenetic analyses revealed that *Hgn. roseipondis* constitutes a novel *Halogranum* species with *Hgn. amylolyticum* and *Hgn. rubrum* being its closest relatives (Fig. 3). The phenotypic properties of *Hgn. roseipondis, Hgn. salarium* (strain of *Hgn. rubrum* used as a reference strain in this study) and *Hgn. amylolyticum* are relatively similar, yet differences in carbon source utilization, gelatin hydrolyzation and genome GC content were observed (Table 1, Fig. 2A). 2A)(12, 14)

Alongside the *Halogranum roseipondis* sp. nov. SS5-1^T^ description, we report here the first characterization of HGTV-1, the only known virus infecting a *Halogranum* species. HGTV-1 displays a unique head-tailed morphology with a long thin neck and wider tail expansion that resembles the classical contractile myovirus tail (Fig. 4). Its lytic infection cycle is relatively slow and modest in burst size (Fig. 4). This might suggest one of two potential scenarios—either it is a finely tuned host-virus interaction possibly shaped by environmental constraints, or HGTV-1 infection proceeds suboptimally in *Hgn. roseipondis* since it is not the preferred host. Although the latter alternative cannot be fully excluded, neither the original cross-infection study (5) nor our subsequent work (Fig. S11) revealed any additional hosts for HGTV-1, as even the closest relative *Hgn. salarium* could not be infected with HGTV-1. A particularly interesting feature of HGTV-1 is its exceptionally large number of viral tRNA genes—covering all amino acids except histidine (Table S3). Codon usage analysis suggests that most of the viral tRNAs may help bridge host-virus codon discrepancies and facilitate the translation of specific viral proteins under host-imposed constraints (Fig. 5). However, not all HGTV-1 encoded tRNAs fit this codon usage hypothesis, possibly implying either a more active role in the translation of host components or other alternative functions. For example, HGTV-1 tRNAs may be differently modified, which could cause ribosomes translating viral transcripts to read through stop codons and/or undergo frameshifting. Some viruses are also known to induce fragmentation of tRNAs (39, 40), yielding specific tRNA fragments that have been reported to suppress antiviral defense mechanisms (41, 42) or directly inhibit translation by interfering with the host translation machinery. Although infection-induced tRNA fragments in archaea have not yet been investigated, stress-induced valine tRNA fragments in *Haloferax volcanii* are known to target the small ribosomal subunit and regulate protein synthesis (43, 44). The continuous evolutionary arms race between hosts and viruses raises the intriguing possibility that viruses may have adapted to exploit also this cellular mechanism. In conclusion, this study provides not only a detailed characterization of a novel virus-host model system, but it establishes a platform for further exploration of the multifaceted mechanisms by which viruses interact with the host translation machinery.

## Materials and methods

### Archaeal strains and virus used in this study

The following archaea were used in this study: *Halogranum roseipondis* sp. *nov*. SS5-1^T^ isolated originally in 2011 (5), and reference strains *Haloarcula vallismortis* (27, 28), *Hgn. salarium* B-1 (13, 26) and *Haloferax gibbonsii* LR2-5 (45). *Har. vallismortis* and *Hgn. salarium* were purchased from the Leibniz Institute DSMZ-German collection of Microorganisms and Cell Cultures GmbH with catalogue numbers DSM 3756 and DSM 23171, respectively. *Hagravirus capitaneum*, HGTV-1, was originally isolated on *Halogranum roseipondis* sp. nov. SS5-1^T^ from a water sample collected from solar saltern in Samut Sakhon, Thailand (5).

### Growth conditions

Modified growth media (MGM) was used for cultivating archaeal strains. The base solution for the medium— a 30% (w/v) stock solution of artificial sea water (SW) containing 4.5 M salts—was prepared as previously published (5) and diluted to 23% (3.4 M salts), 20% (3.0 M salts) and 18% (2.7 M salts) (v/v) for liquid culture, solid and top-layer media, respectively. Solid and top-layer media were obtained by adding 14 g or 4 g of Bacto agar (Difco Laboratories) per liter, respectively, and included 5 g peptone (Oxoid) and 1 g Bacto yeast extract (Difco Laboratories) per liter. For assessing growth at different salt conditions, the dilution of SW in the media was adjusted to obtain the desired final salt concentration. Archaeal strains were streaked on MGM-agar plates and grown at 37 °C until colonies appeared. The liquid starter cultures were inoculated in 30 ml of MGM broth and grown for 72 h at 37 °C with 200 rpm aeration. For growth temperature characterization, the starter culture was grown at the corresponding temperature.

The minimum NaCl requirement to maintain cells intact was tested by harvesting cells from 0.5 ml volume of a stater culture by centrifugation [(3000x*g*, 5 min, room temperature (RT)] and gently resuspending the cells to NaCl in ddH_2_O (0-18% NaCl range). Cells were washed by pelleting and gently resuspending them to the desired NaCl concentration. Cell integrity was observed after 30 min and 15 h of incubation at RT using phase contrast microscopy with Leica DM750 microscope.

Optimal culture conditions were determined using a Bioscreen FP-1100-C by assessing growth (defined as change in optical density at 600 nm (OD_600_)) at various temperatures, pH, and salinity ranges. Briefly, starter cultures were diluted to OD_600_ 0.1-0.2 using fresh medium (corresponding to the condition tested) and loaded onto Bioscreen honeycomb well plates. Alternatively, starter cultures were not diluted but used as such if OD_600_ <0.1. All growth curve characterizations were performed with a total culture volume of 380 µL/well. The Bioscreen was set to continuous medium aeration and amplitude with temperature control applied according to the condition tested.

### Genome isolation and analysis

The gDNA was isolated by spooling as previously described (46) with minor modifications. Briefly, the 50 mL of exponential culture (OD_600_ = 0.6) was pelleted by centrifugation at 8 000x*g*, 10 min, 15 °C. The cell pellet was resuspended in 2 mL ST buffer (1M NaCl, 20 mM Tris-HCl, pH 7.5). Cell suspension was lysed by addition of 2 mL Lysis solution (100 mM EDTA pH 8.0, 0.2% SDS). The mixture was overlayed with 4 mL of 99.6% ethanol and gDNA was spooled from the interphase by capillary. The spooled gDNA was washed twice by swirling the capillary in 1 mL of 99.6% ethanol. The gDNA was then dried, resuspended in 1 mL TE buffer and finally precipitated by addition of 0.1 vol of 3 M sodium acetate pH 5.5 and 0.8 vol of isopropanol. After pelleting (6 000x*g*, 5 min, RT), the gDNA was washed by 1 mL of 70% ethanol and pelleted again. The air-dried pellet was resuspended in 1 mL TE and treated by 1 µl (30 mg/mL) DNase-free RNase A (Applichem) for 1h at 45 °C. The gDNA was re-isolated by isopropanol precipitation as described above and the final pellet was resuspended in TE.

The genome was sequenced using Pacific Biosciences PacBio Sequel II sequencing technology and assembly was performed using PacBio Microbial Genome Analysis application integrated in SMRTlink v11 at the DNA Sequencing and Genomics Laboratory, Helsinki Institute of Life Science, University of Helsinki. The assembled genome was annotated using prokaryotic genome annotation tool Prokka (47), Galaxy version 1.14.6+galaxy1. DNA molecules were visualized using DNAplotter (48). The genome sequence has been submitted to GenBank under the temporary submission ID: SUB15737247.

The phylogeny of *Hgn. roseipondis* was first investigated by pangenome analysis using Roary (49) and further assessed by comparing 16S rRNA sequences of all *Halogranum* species and the reference strains using MUltiple Sequence Comparison by Log-Expectation (MUSCLE) with default settings (50) and by conducting average nucleotide identity by orthology, OrthoANI (51). Roary and OrthoANI analyses were applied on a set of 125 type species representing all genera within family *Haloferacaceae* except *Candidatus Haloaextosymbiotes* (Table S1).

HGTV-1 genome was retrieved from NCBI GenBank with accession number NC_021328. tRNAscan-SE 2.0 (32) was used in default mode with sequence sources for archaea or mixed settings were selected to find tRNA genes in *Hgn. roseipondis* and HGTV-1 genomes, respectively. Codon usage was analyzed with Codon usage (Cusp) application from the European Molecular Biology Open Software Suite (52). Sequence comparison of host and virus tRNAs was conducted using MUSCLE (50) with default settings.

### Phenotypic assays

For subsequent assays, *Hgn. roseinpondis* grown on an MGM plate for 3 to 4 nights was used. Colonies were streaked on a glass slide and supplemented with a droplet of ∼40 µl of 3% H_2_O_2_. Catalase activity was indicated by the release of oxygen, observed as bubbling within the culture mass. The oxidase test was performed by applying 1% N, N, N′, N′-tetramethyl-p-phenylenediamine dihydrochloride (in ddH_2_O) on a cotton swab with archaeal mass from the plate and the oxidase activity was detected by the formation of dark purple color.

Next, single carbon source utilization was determined as described in the Halohandbook (46, 53). Briefly, minimal media including 1 ml/L of trace element solution (2.5 mM MnCl_2_, 2.2 mM ZnSO_4_, 0.2 mM CuSO_4_, 8.3 mM FeSO_4_) was prepared and supplemented with glycerol (0.3% w/v), sodium acetate (0.1% w/v), sodium succinate (0.5% w/v), galactose (0.4% w/v) or 0.3% w/v fructose, glucose, lactose, mannitol, or sucrose. Three separate colonies were streaked on a minimal media plate and growth was assessed after a 14-day incubation at 37 °C.

Inhibition of the growth by different antibiotics was tested by growing the archaea in 23% MGM liquid broth supplemented with ampicillin 100 µg/mL, streptomycin 50 µg/mL, tetracycline 10 µg/mL, chloramphenicol 35 µg/mL, rifampicin 10 µg/mL, novobiocin 2.5 µg/mL or mevinolin 4 µg/mL.

Anaerobic growth was assessed by streaking MGM plates and incubating them in an anaerobic jar, from which oxygen was depleted using Microbiology Anaerocult A (Merck) strips according to the manufacturer’s instructions. Potential growth was assessed following a 7-day incubation period.

For starch hydrolysis, archaeal strains were streaked onto 20% MGM agar plates supplemented with soluble starch (2 g/L) and incubated for one week at 37 °C. Hydrolysis was detected by flooding the plates with Lugol’s iodine solution (54). Gelatin hydrolysis tests were performed by utilizing 20% MGM agar plates supplemented with 0.8% (w/v) gelatin, according to the protocol by Dela Cruz and Torres (55). After a 14-day incubation at 37 °C, results were visualized by flooding the plate with saturated ammonium sulphate solution.

Motility was assessed as spread of growth on top layer soft-agar plates by either stabbing colonies into the top-layer agar or by pipetting 10 µL droplets of the early exponential phase liquid culture on the plates and by observing the cells using light microscopy. Plates were incubated at 37°C for 5 nights. For light microscopy, samples from plate and liquid culture were prepared. Colonies were mixed in 20% SW and liquid culture grown to OD_600_ of 0.6 was directly pipetted on microscopy slide.

### Electron microscopy

Transmission electron microscopy (TEM) and scanning electron microscopy (SEM) of *Hgn. roseipondis* cells was conducted using liquid cultures grown to OD_600_ 0.6 (early) or 1.0 (mid-logarithmic), respectively. 5 mL of cells were collected by centrifugation at 3400x*g* for 5 min, resuspended in 23% SW and fixed in 2.5% glutaraldehyde at 4°C for 18 h. Glutaraldehyde was removed by centrifugation as above and replaced with water. For TEM, 2 µL of the fixed cell sample was pipetted on a carbon coated, glow discharged copper mesh grid for 2 min, after which the sample was negative stained using 3% uranyl acetate for 10 s and rinsed with ddH_2_O. For SEM, fixed cells were immobilized on concavalin A coated glass slips, osmicated with 1% OsO_4_ (in water) and dehydrated through ethanol dehydration series with 50% (v/v), 70%, 96% and 99.6% ethanol. Microscopy was performed using

Virus sample for TEM was produced by purifying the virus from a liquid culture infected at OD_600_ of 0.6 as has been described before (5). In short, viruses were precipitated from the lysate using polyethylene glycol 6000 (10% w/v) and further purified through rate-zonal ultracentrifugation in linear 5-20% sucrose gradient prepared in 18% SW (104 000x*g*, 30 min, 20 °C). Light scattering virus zone from the gradient was collected and viruses were concentrated using differential ultracentrifugation (128 000x*g*, 75 min, 15 °C). The purified virus was resuspended in 18% SW, pipetted on a glow discharged, carbon coated Cu-grid and stained with neutral 2% uranyl acetate.

TEM was performed with JEOL JEM-1400 (Jeol Ltd., Tokyo, Japan) equipped with Gatan Orius SC 1000B CCD-camera (Gatan Inc. USA) and SEM with a FEI Quanta 250 Field Emission Gun scanning electron microscope equipped with Gatan 3View system. Electron Microscopy was carried out in Electron Microscopy Unit of the Institute of Biotechnology, University of Helsinki.

### HGTV-1 infection analysis

HGTV-1 plate lysate was prepared by plating suitably diluted virus yielding semiconfluent plates mixed with 200 µl of host starter culture and 3 mL of 18% MGM top-agar (temperature adjusted to 55 °C) on 20% MGM agar-plates, which were incubated at 37°C for 4 nights. The top-agar layer from each semiconfluent plate was collected to a flask, to which 1.8 mL of 23% MGM broth/plate was added. Next, viruses were separated from agar and cell debris by shaking the mixture at RT for 2 h, 200 rpm, followed by centrifugation at 8800x*g* for 30 min at RT.

To estimate the number of infective viruses, plaque assays were performed for various samples throughout this study. Briefly, a dilution series of the virus sample is made in 23% MGM to yield a countable number of plaques on the host lawn and plated as described above. The average of plaques produced from each dilution is then applied to count the number of infective viruses in the original sample.

The cross-infectivity of HGTV-1 was tested also on the reference strains (see section “Archaeal strains and virus used in this study”) by conducting a spot-on-lawn assay. The archaeal starter culture was plated with top-agar as described above and let to solidify for ∼30 min at RT. Different dilutions of HGTV-1 plate lysate was pipetted on each plate as 10 µL droplets and the infectivity was observed as plaque formation after incubation of the plates in 37 °C for 5 nights.

HGTV-1 infection on *Hgn. roseipondis* was investigated in different salinities and temperatures. *Hgn. roseipondis* liquid cultures were grown to OD_600_ 0.6 in growth media of different salinities and temperatures and infected with plate lysate using multiplicity of infection (MOI) 10. Cultures were then divided on honeycomb plates and optical density of the cultures were monitored using Bioscreen FP-1100-C as described earlier.

Length of the intracellular phase of the infection cycle and the burst size of HGTV-1 was determined by conducting a one-step growth curve assay (56) in which the applied adsorption time with light shaking after adding the viruses to the culture was 3 h.

## Acknowledgements

The authors are grateful to Dr Hanna Oksanen for providing the *Hgn. roseinpondis* sp. nov. SS5-1^T^ from the Samut Sakhon collection initiated by Prof. Em. Dennis Bamford and the *Hfx. gibbonsii* LR2-5 strain. The authors wish to thank the following University of Helsinki core facilities: Electron Microscopy Unit, Institute of Biotechnology, HiLIFE—SEM sample preparation, instrument access, and user support; Instruct-HiLIFE Biocomplex, a member of Instruct-ERIC Centre Finland, FINStruct, and Biocenter Finland—ultracentrifugation services; and DNA Sequencing and Genomics, Institute of Biotechnology, HiLIFE—whole genome sequencing services. CSC—IT Center for Science Ltd., Finland, is acknowledged for computational resources. All members of the RNAcious laboratory are warmly acknowledged for insightful discussions. P.G. is a fellow of the Doctoral Programme in Integrative Life Science, University of Helsinki Doctoral School.

This publication is dedicated to the memory of Prof. Em. Dennis H. Bamford (1948-2025).

## Funding

This research was funded by the Novo Nordisk Foundation (Postdoctoral Fellowships for research within Industrial Biotechnology and Environmental Biotechnology), grant no. NNF22OC0079323, to M.J.L.; Emerging Investigator in Biotechnology-Based Synthesis and Production, grant no. NNF19OC0054454, to L.P.S.) and Research Council of Finland (Academy Project, grant no. 354906, to L.P.S.).

## Notes

### Competing Interest Statement

The authors have declared no competing interest.

