## Supplemental data for "The halophilic archaeon *Halogranum roseipondis* sp. nov. is susceptible to a virus carrying an exceptionally high number of viral tRNA genes"

**Contents**

Fig. S1. *Halogranum roseipondis* SS5-1$^T$ is non-motile in laboratory conditions.

Fig. S2. *Halogranum roseipondis* SS5-1$^T$ pigmentation is weaker at high temperatures.

Fig. S3. Liquid microcultures of *Hgn. roseipondis* SS5-1$^T$ and reference strains grown at different salinity and temperature conditions.

Fig. S4. Liquid microcultures of *Hgn. roseipondis* SS5-1$^T$ grown at different pH conditions.

Fig. S5. *Hgn. roseipondis* SS5-1$^T$ hydrolyzes starch but not gelatin

Fig. S6. The effect of different antibiotics on the growth of *Hgn. roseipondis* SS5-1$^T$ and reference strains.

Fig. S7. Transmission electron microscopy on *Halogranum roseipondis* SS5-1$^T$ reveals irregular cell morphology

Fig. S8. Roary pangenomic analysis of representative type species from all genera within family *Haloferacaceae*

Fig. S9. OrthoANI analysis of representative type species from all genera within family *Haloferacaceae*

Fig. S10. Detached HGTV-1 tails and possible non-processed tail assembly intermediates

Fig. S11. HGTV-1 infection on different hosts

Fig. S12. HGTV-1 infection in different temperatures and salinities

Fig. S13. Comparison of tRNA sequence similarity for a selection of *Hgn. roseipondis* SS5-1$^T$ and HGTV1 tRNAs.

Table S1. Sequencing depth

Table S2. Strains and their accession numbers applied in the Roary and ANI analysis

Table S3. HGTV-1 tRNAs

Table S4. Codon usage in *Halogranum roseipondis* SS5-1$^T$ and HGTV-1 genomes

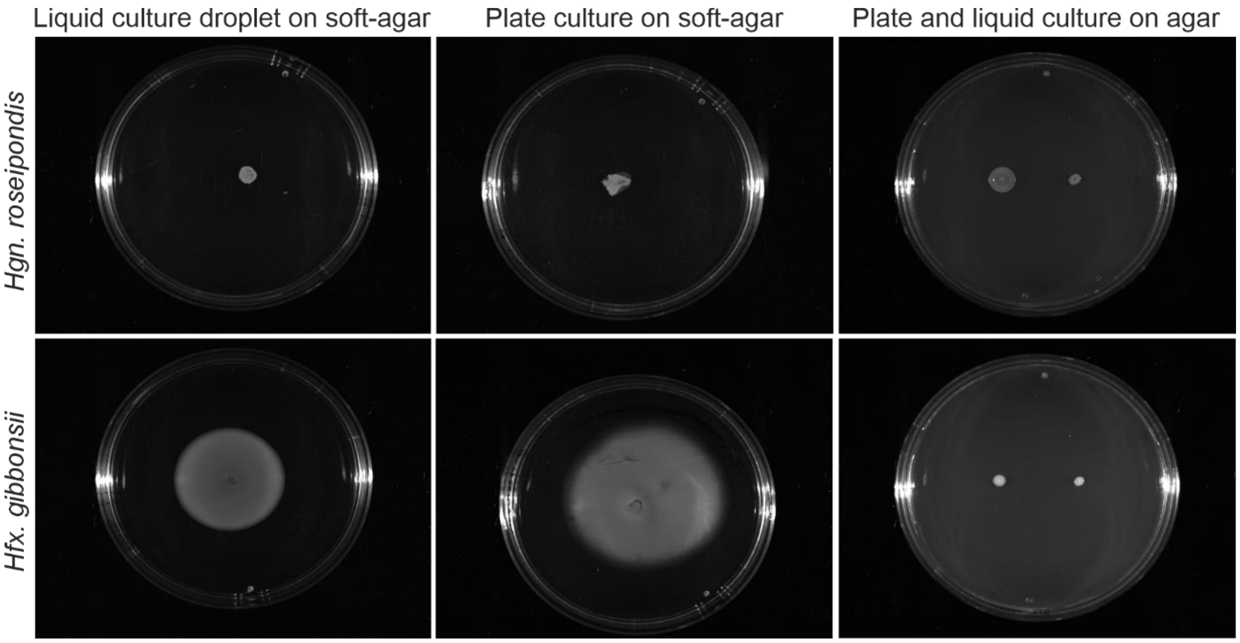


**FIG S1. *Halogranum roseipondis* SS5-1$^T$ is non-motile in laboratory conditions.** Motility assay of *Hgn. roseipondis* from liquid and plate cultures grown on soft-agar and solid-agar plates (top panels). *Hfx. gibbonsii* cultures are included for comparison (bottom panels).

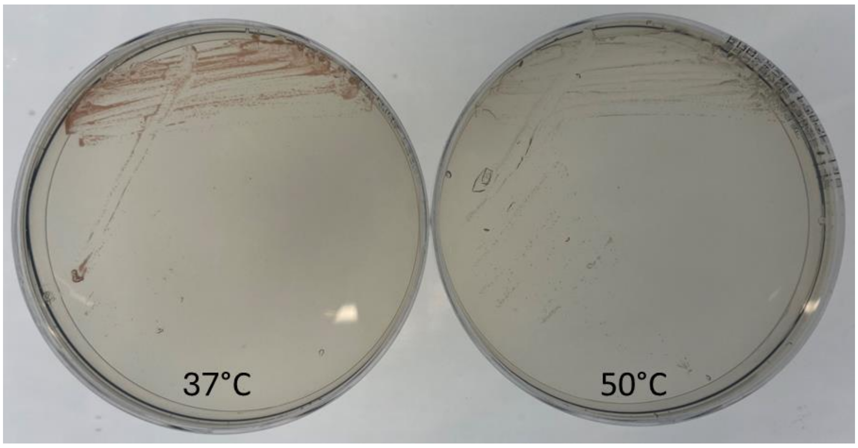


**FIG S2. *Halogranum roseipondis* SS5-1$^T$ pigmentation is weaker at high temperatures.** Left: *Hgn. roseipondis* grown on a 20% MGM plate at 37 °C. Right: *Hgn. roseipondis* grown on a 20% MGM plate at 50 °C. Note the lack of pink/reddish coloration at 50 °C.

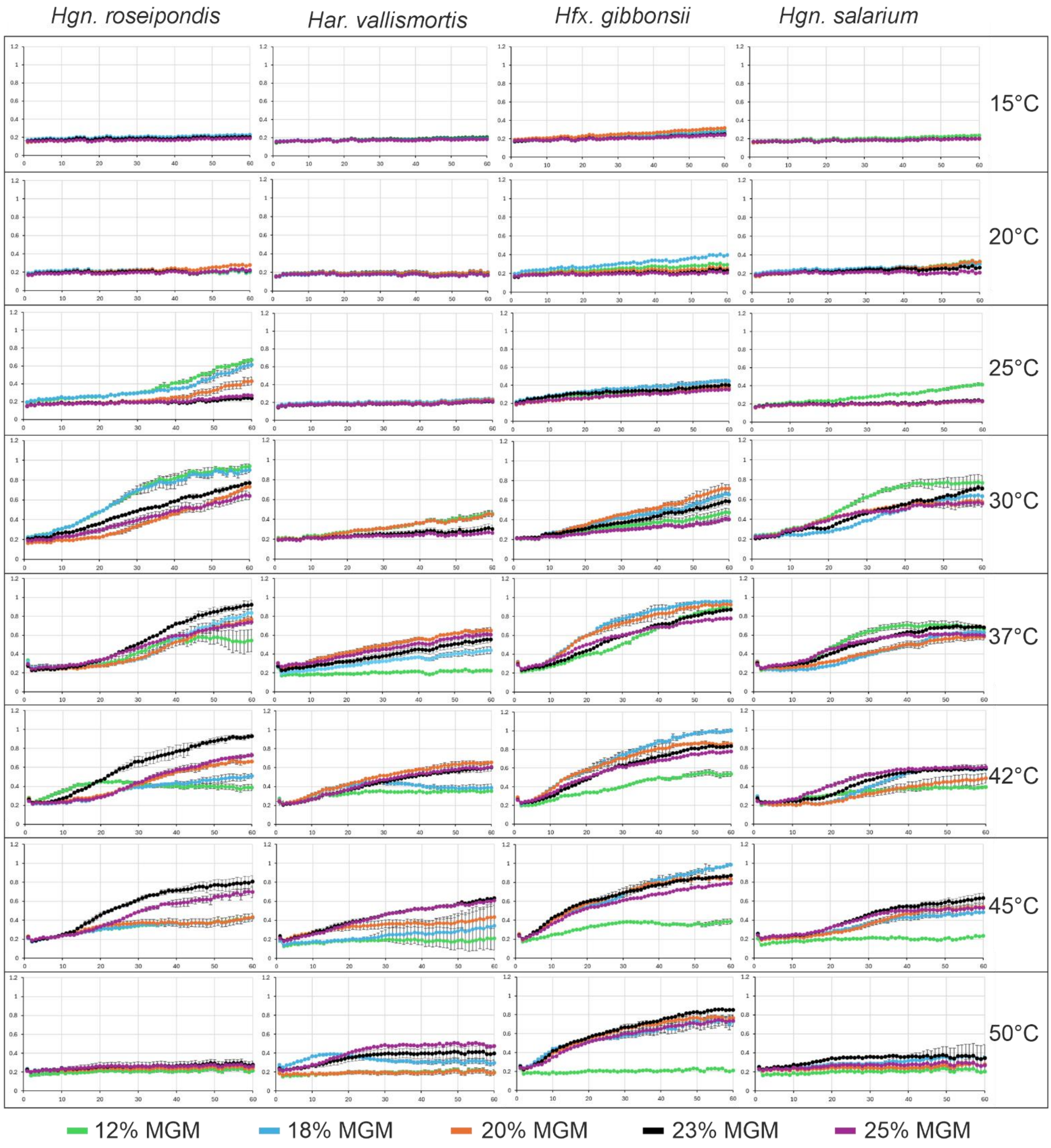


**FIG S3. Liquid microcultures of *Hgn. roseipondis* SS5-1[T] and reference strains grown at different salinity and temperature conditions.** Growth curves plotted as $OD_{600}$ (y-axis, 0-1.2) against time (x-axis, 0-60 h). Error bars indicate standard deviation (n=5).

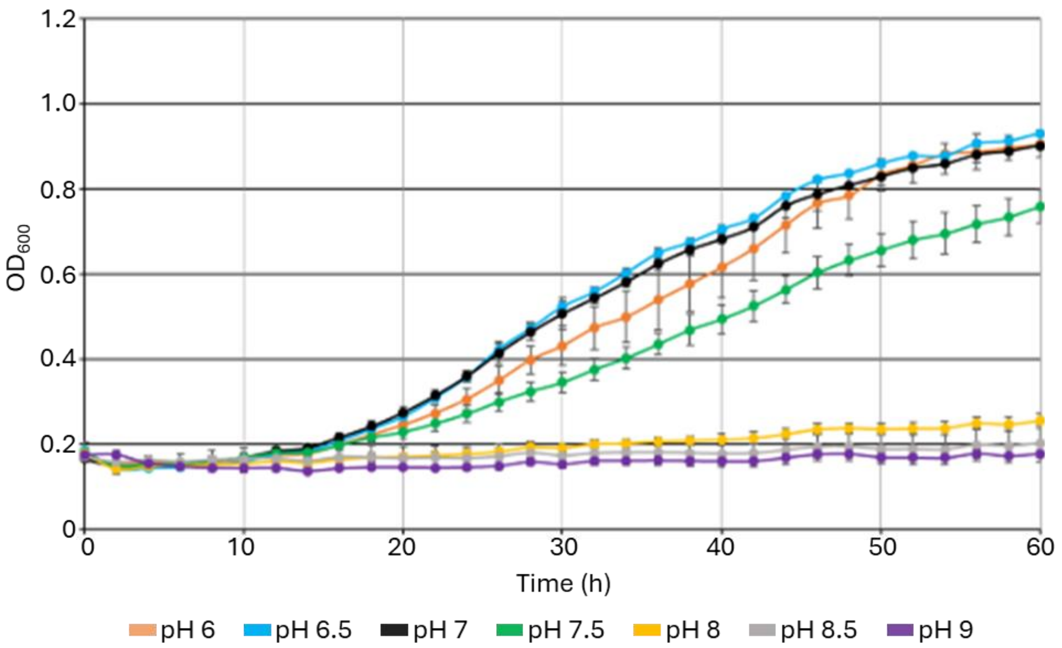


**FIG S4. Liquid microcultures of *Hgn. roseipondis* SS5-1$^T$ grown at different pH conditions.** Error bars indicate standard deviation (n=5).

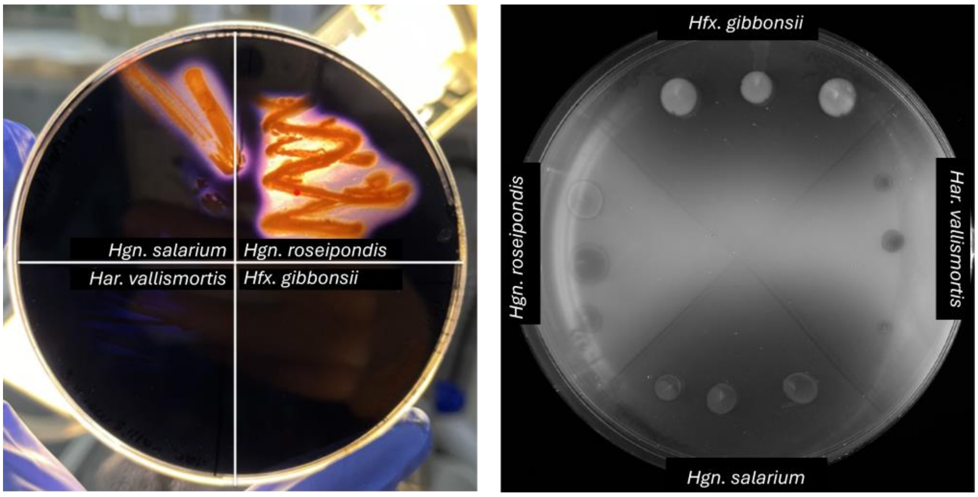


**FIG S5. *Hgn. roseipondis* SS5-1$^T$ hydrolyzes starch but not gelatin**. Starch hydrolyzation by *Hgn. roseipondis* and *Hgn. salarium* was observed as bright halos on starch-MGM plate after flooding the plate with Lugol's iodine solution (left). Gelatin hydrolyzation on a gelatin-MGM plate flooded with saturated ammonium sulphate was not observed for *Hgn. roseipondis*, whereas clear halos were observed for *Hgn. salarium* and *Hfx. gibbonsii*, respectively (right).

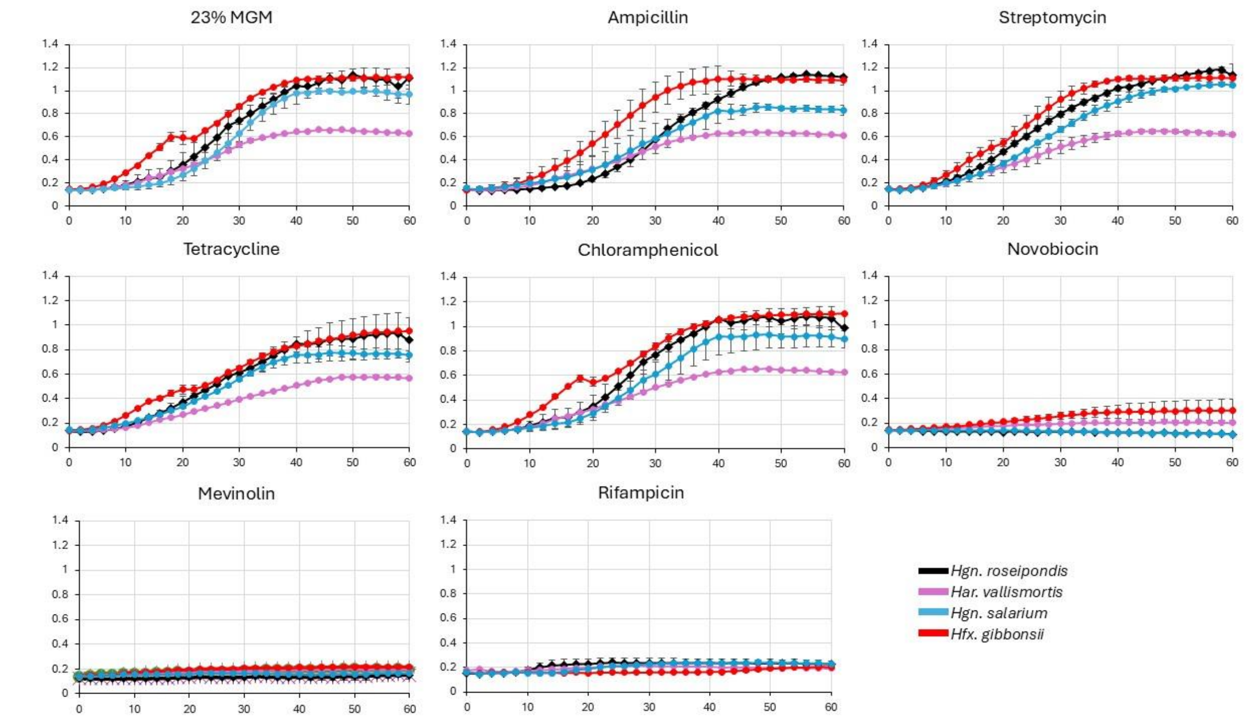


**FIG S6. The effect of different antibiotics on the growth of *Hgn. roseipondis* SS5-1$^T$ and reference strains.** Growth curves plotted as $OD_{600}$ (y-axis, 0-1.4) against time (x-axis, 0-60 h). Error bars indicate standard deviation (n=3).

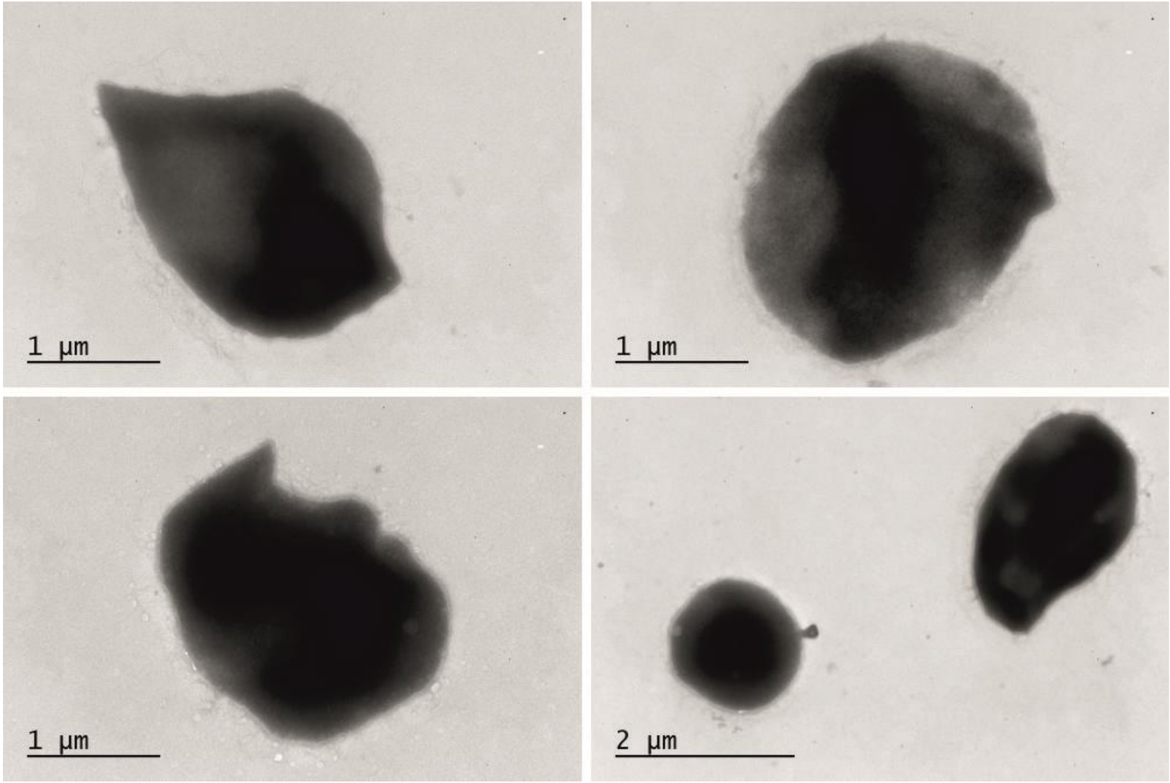


**FIG S7. Transmission electron micrographs of *Halogranum roseipondis* SS5-1$^T$ reveals irregular cell morphology.**

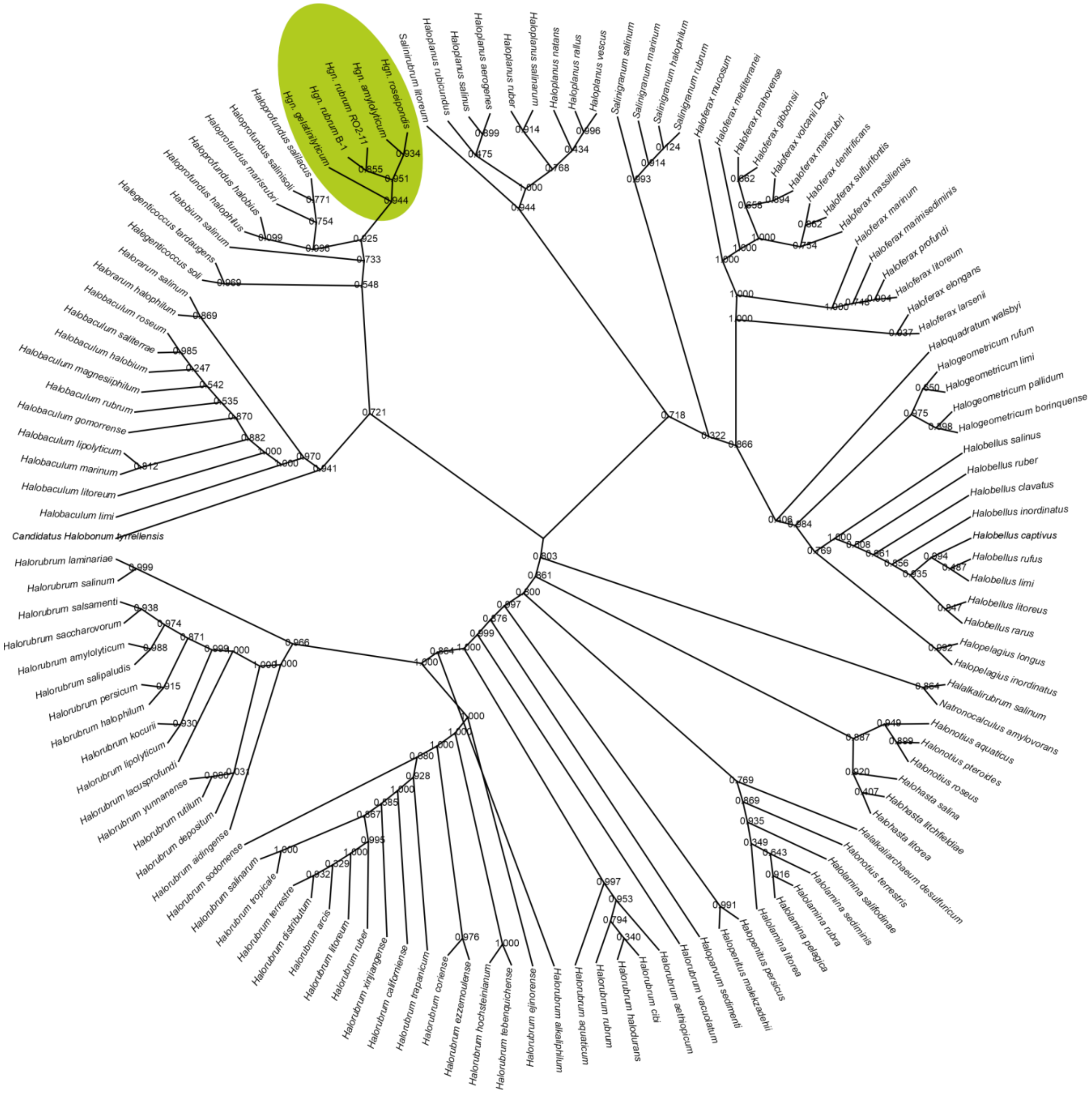


**FIG S8. Roary pangenomic analysis of representative type species from all genera within family *Haloferacaceae*.** Genus *Halogranum* highlighted with green background.

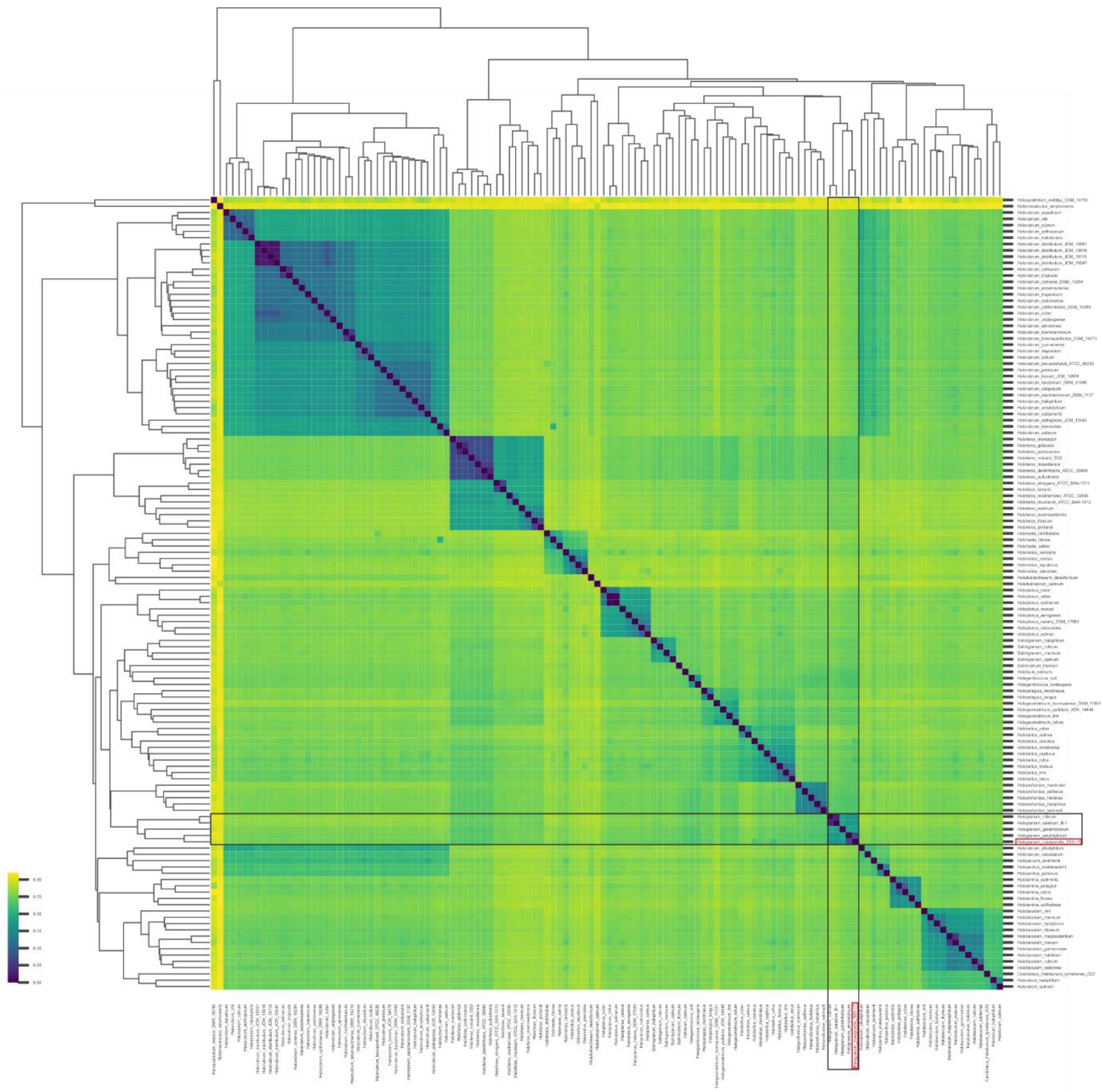

**FIG S9. OrthoANI analysis of representative type species from all genera within the family *Haloferacaceae*.** Species of *Halogranum* genus are marked with black boxes. In addition, *Halogranum roseipondis* SS5-1$^{T}$ is indicated with red boxes.

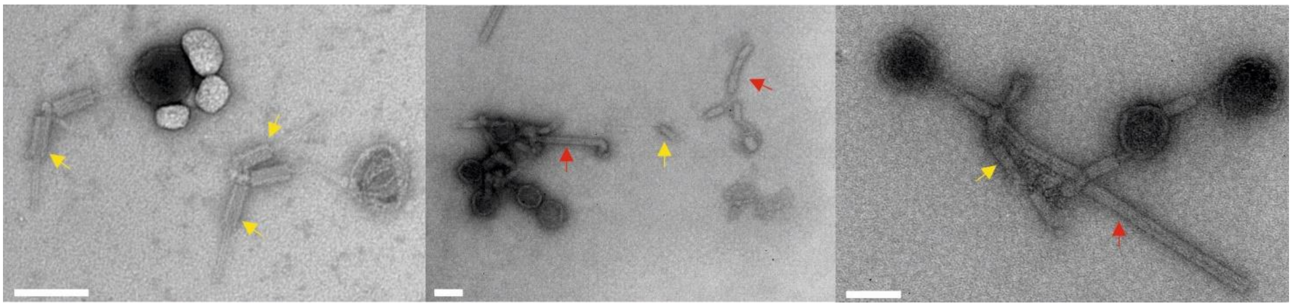

**FIG S10. Detached HGTV-1 tails and possible non-processed tail assembly intermediates.** Yellow arrows indicate detached/loose tail structures and red arrows point to possible tail assembly intermediates. Size bar is 100 nm in each panel

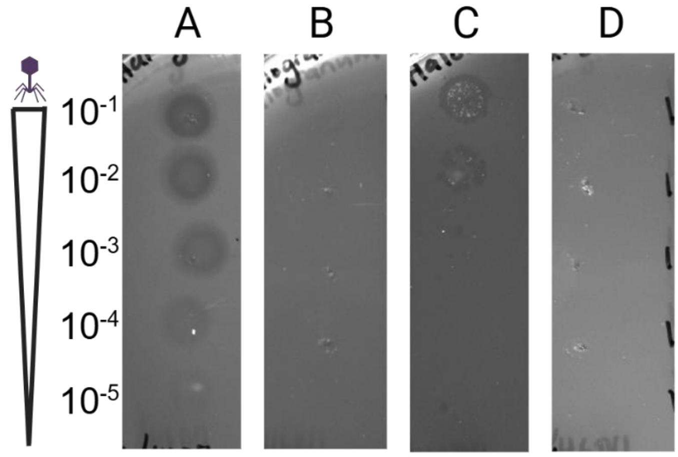


**FIG S11. HGTV-1 infection on different hosts.** A) Strong HGTV-1 infection on the isolation host *Hgn. roseipondis* SS5-1$^{T}$ is observed. B) HGTV-1 is not infective on *Hgn. salarium*, C) Weak HGTV-1 infection is detected on *Har. vallismortis,* and D) HGTV-1 is not infective on *Hfx. gibbonsii*. Dilutions from $10^{-1}$ to $10^{-5}$ (400000-4 pfu/droplet) of the virus stock solution were applied for testing. The corresponding dilutions are indicated on right.

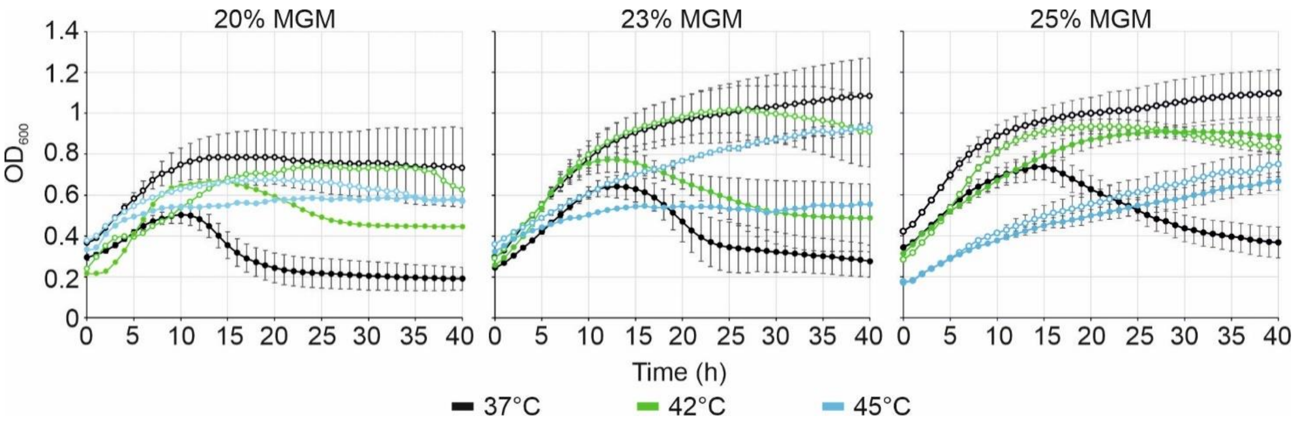


**FIG S12. HGTV-1 infection curves at different temperatures and salinities.** The curves with empty marker circles are mock-infected cultures and the curves with filled marker circles display the infected cultures. Error bars depict the standard deviation (n=3)

**Alignment**

```
HGTV_trna7_Glu        -----GCTGATGGAGTCTCCGCTGGCGAGAGACGCTACGTTCTCAGCGTAGAGGTCGCGGGTTCAATTCCTGTCATCAGCA
Hgr_chr_trna23_Glu    GCTCGGTTGGTGTAGT-----CCGGCCAATCATCTTGGCCTTTC-GAGCCGAGGACAGGGGTTCAAATCCCCTACCGAGCA
Hgr_chr_trna25_Glu    GCTCTGTTGGTGTAGT-----CCGGCCAATCATATCACCCTCTC-ACGGTGATGACCAGGGTTCGAATCCCTGACGGAGCA
```

**Percent identity matrix**

| | | | |
|---|---|---|---|
| HGTV_trna7_Glu | 100.00 | 58.57 | 57.14 |
| Hgr_chr_trna23_Glu | 58.57 | 100.00 | 77.33 |
| Hgr_chr_trna25_Glu | 57.14 | 77.33 | 100.00 |

**Phylogram**

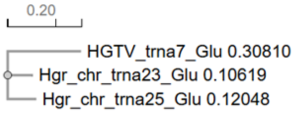


**Alignment**

```
Hgr_chr_trna41_Gln    GAAAGGGTCGCTCAGCCTGGCCAGAGCGCCCACCATTGCGGTGGGTGGCCTGAAAGATGGCCACGATGGTTCAAATCCATCCCCTTTCG
HGTV_trna19_Gln       ----AGTCCCCGAGTCCGCT-----GGCGAAAGACGCTCGGCTTTGGACCGAGAGAGCGA------GGTTCGATTCCTCGGGGGACTA
Hgr_chr_trna10_Gln    -----AGTTCCATGGGGTAGT-----GGCCAATCCTGTTGCCTTCTGGGGGCAACGACCCA------GGTTCGAATCCTGGTGGAACTA
HGTV_trna23_Gln       -----AGTCGGTTTGTGTAGC-----GGCCAATCATACGAGGCTCTGACCCTCGTGACAGA------GGTTCGAATCCTCTACCGACTA
Hgr_chr.trna55_Gln    -----AGTCCCGTGGTGTAGT-----GGCCAATCATCTGGGCCTTTGGAGCCCAGGACAGC------GGTTCGAATCCGCTCGGGACTA
```

**Percent identity matrix**

| | | | | | |
|---|---|---|---|---|---|
| Hgr_chr_trna41_Gln | 100.00 | 43.84 | 41.10 | 49.32 | 47.95 |
| HGTV_trna19_Gln | 43.84 | 100.00 | 56.16 | 58.90 | 61.64 |
| Hgr_chr_trna10_Gln | 41.10 | 56.16 | 100.00 | 57.53 | 71.23 |
| HGTV_trna23_Gln | 49.32 | 58.90 | 57.53 | 100.00 | 71.23 |
| Hgr_chr.trna55_Gln | 47.95 | 61.64 | 71.23 | 71.23 | 100.00 |

**Phylogram**

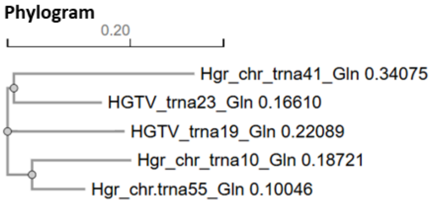


**Alignment**

```
HGTV_trna8_Gly        GCGTGGCTGTTCCAATGGCAAGATGCGGGCCTTCCAAGCCTGAGATACGAGTTCGACTCTCGTGCCGCGCA
HGTV_trna9_Gly        GCGTGGTTGGTCCAATGGAAAGACGGCTCCCTGCCACGGAGCAGACTCGGGTTCGATTCCCGAATCGCGCA
Hgr_chr_trna38_Gly    GCACTGATAGTGTAGTGGTATCACGTGACCTTGCCATGGTCACAACCTGGGTTCAAATCCCAGTCAGTGCA
Hgr_chr_trna39_Gly    GCACTGATAGTGTAGTGGTATCACGTGACCTTGCCATGGTCACAACCTGGGTTCAAATCCCAGTCAGTGCA
Hgr_chr_trna56_Gly    GCACCGGTGGTCTAATGGCATGACTTTGGCCTTCCAAGCCAACGATCTGGGTTCAACTCCCGGCCGGTGCA
Hgr_chr_trna14_Gly    GCGCCGTTGGTCCAGTGGTAGGACAGTGGCTTCCCAAGCCACTAGCCCGGGTTCAATTCCCGGACGGCGCA
```

**Percent identity matrix**

| | | | | | | |
|---|---|---|---|---|---|---|
| HGTV_trna8_Gly | 100.00 | 69.01 | 47.89 | 47.89 | 67.61 | 61.97 |
| HGTV_trna9_Gly | 69.01 | 100.00 | 56.34 | 56.34 | 59.15 | 67.61 |
| Hgr_chr_trna38_Gly | 47.89 | 56.34 | 100.00 | 100.00 | 67.61 | 61.97 |
| Hgr_chr_trna39_Gly | 47.89 | 56.34 | 100.00 | 100.00 | 67.61 | 61.97 |
| Hgr_chr_trna56_Gly | 67.61 | 59.15 | 67.61 | 67.61 | 100.00 | 73.24 |
| Hgr_chr_trna14_Gly | 61.97 | 67.61 | 61.97 | 61.97 | 73.24 | 100.00 |

**Phylogram**

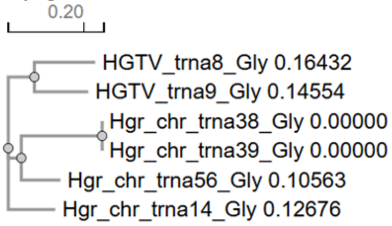


**Alignment**

```
HGTV_trna24_Ile       GCGTTCGTAGCCAAATCAGGTA-AGGCGTTCCGCTGATAACGGAAAGATTTGTTGGTTCAAATCCAACCGGACGCA
Hgr_chr_trna8_Ile2    GGGCCCTTAGCTCAGTCTGGTTAGAGCGCTCGGCTCATAACCGAGTGGTCAT-TGGTTCGAATCCGATAGGGCCCA
Hgr_chr_trna35_Ile    GGGCCAATAGCTCAGTCAGGTT-GAGCGCTCGGCTGATAACCGGGAGGTCCG-CGGTTCAAATCCGCGTTGGCCCA
```

**Percent identity matrix**

| | | | |
|---|---|---|---|
| HGTV_trna24_Ile | 100.00 | 63.51 | 63.51 |
| Hgr_chr_trna8_Ile2 | 63.51 | 100.00 | 81.08 |
| Hgr_chr_trna35_Ile | 63.51 | 81.08 | 100.00 |

**Phylogram**

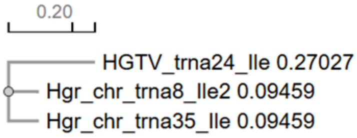


**FIG S13.** Comparison of tRNA sequence similarity for a selection of *Hgn. roseipondis* SS5-1$^{T}$ and HGTV1 tRNAs. Alignment of tRNA sequences, percent identity matrix and its visualization as a phylogram. Naming of the tRNAs correspond to Supplementary Table 3.

### Table S1. PacBio HiFi sequencing depth

| Molecule | Average coverage depth |
|---|---|
| **Hgr_chr** | 18.7 |
| **Hgr_p560** | 23.3 |
| **Hgr_p325** | 36.7 |
| **Hgr_p222** | 29.4 |
| **Hgr_p194** | 40.2 |
| **Hgr_p151** | 34.0 |
| **Hgr_63** | 34.9 |
| **Hgr_p40** | 44.2 |

### Table S2. Reference genomes used for Roary pangenomic analysis

| Strain | Accession | Assembly |
|---|---|---|
| *Halogranum roseipondis* SS5-1T | | chromosome and 7 plasmids |
| *Halogranum amylolyticum* | NZ_FODV01000001.1 - NZ_FODV01000061.1 | scaffold |
| *Halogranum rubrum* R02-11 | NZ_FOTC01000001.1 - NZ_FOTC01000017.1 | contig |
| *Halogranum rubrum* B-1 | NZ_ALJD01000001.1 - NZ_ALJD01000017.1 | contig |
| *Halogranum gelatinilyticum* | NZ_FNHL01000001.1 - NZ_FNHL01000011.1 | contig |
| *Haloprofundus salilacus* | NZ_CP083723.1, NZ_CP083724.1 | two chromosomes |
| *Haloprofundus salinisoli* | NZ_CP083663.1, NZ_CP083664.1, NZ_CP083665.1 | chromosome, two plasmids |
| *Haloprofundus marisrubri* | NZ_LOPU01000001.1 - NZ_LOPU01000041.1 | contig |
| *Haloprofundus halobius* | NZ_CP083666.1, NZ_CP083667.1 - NZ_CP083670.1 | chromosome, four plasmids |
| *Haloprofundus halophilus* | NZ_QQRR01000001.1 - NZ_QQRR01000006.1 | scaffold |
| *Halobium salinum* | NZ_JAODIW010000001.1 - NZ_JAODIW010000014.1 | scaffold |
| *Halegenticoccus tardaugens* | NZ_SDIC01000001.1 - NZ_SDIC01000014.1 | contig |
| *Halegenticoccus soli* | NZ_PEND01000001.1 - NZ_PEND01000014.1, NZ_KZ537954.1 | contig |
| *Halorarum salinum* | NZ_CP058579.1, NZ_CP058580.1 | chromosome, plasmid |
| *Halorarum halophilum* | NZ_CP058529.1, NZ_CP058530.1 - NZ_CP058532.1 | chromosome, three plasmids |
| *Halobaculum roseum* | NZ_CP082286.1, NZ_CP082287.1 - NZ_CP082289.1 | chromosome, three plasmids |
| *Halobaculum saliterrae* | NZ_WUUS01000001.1 - NZ_WUUS01000021.1 | contig |
| *Halobaculum halobium* | NZ_CP126158.1 - NZ_CP126160.1 | chromosome, two plasmids |
| *Halobaculum magnesiiphilum* | NZ_CP081958.1, NZ_CP081959.1 - NZ_CP081961.1 | chromosome, three plasmids |
| *Halobaculum rubrum* | NZ_CP082284.1, NZ_CP082285.1 | chromosome, plasmid |
| *Halobaculum gomorrense* | NZ_FQWV01000001.1 - NZ_FQWV01000016.1 | scaffold |
| *Halobaculum lipolyticum* | NZ_CP126154.1, NZ_CP126155.1 | chromosome, plasmid |

| | | |
|---|---|---|
| *Halobaculum marinum* | NZ_CP119989.1, NZ_CP119990.1, NZ_CP119991.1 | chromosome, two plasmids |
| *Halobaculum litoreum* | NZ_CP126156.1, NZ_CP126157.1 | chromosome, plasmid |
| *Halobaculum limi* | NZ_CP120468.1, NZ_CP120469.1 | chromosome, plasmid |
| *Candidatus Halobonum tyrrellensis* | NZ_ASGZ01000001.1 - NZ_ASGZ01000072.1 | contig |
| *Halorubrum laminariae* | NZ_JANHDL010000001.1 - NZ_JANHDL010000042.1 | scaffold |
| *Halorubrum salinum* | NZ_JANHDO010000001.1 - NZ_JANHDO010000006.1 | scaffold |
| *Halorubrum salsamenti* | NZ_VCNL01000001.1 - NZ_VCNL01000021.1 | scaffold |
| *Halorubrum saccharovorum* | NZ_AOJE01000001.1 - NZ_AOJE01000072.1 | contig |
| *Halorubrum amylolyticum* | NZ_SDJP01000001.1 - NZ_SDJP01000075.1 | scaffold |
| *Halorubrum salipaludis* | NZ_NSKC01000001.1 - NZ_NSKC01000027.1 | contig |
| *Halorubrum persicum* | NZ_NHOA01000001.1 - NZ_NHOA01000158.1 | scaffold |
| *Halorubrum halophilum* | NZ_BBJP01000001.1 - NZ_BBJP01000103.1 | contig |
| *Halorubrum kocurii* | NZ_AOJH01000001.1 - NZ_AOJH01000105.1 | contig |
| *Halorubrum lipolyticum* | NZ_AOJG01000001.1 - NZ_AOJG01000041.1 | contig |
| *Halorubrum lacusprofundi* | NC_012028.1, NC_012029.1, NC_012030.1 | chromosomes 1 and 2, plasmid pHLAC01 |
| *Halorubrum yunnanense* | NZ_JAODIX010000001.1 - NZ_JAODIX010000116.1 | scaffold |
| *Halorubrum rutilum* | NZ_JANHDN010000001.1 - NZ_JANHDN010000012.1 | scaffold |
| *Halorubrum depositum* | NZ_VCNM01000001.1 - NZ_VCNM01000005.1 | scaffold |
| *Halorubrum aidigense* | NZ_AOJI01000001.1 - NZ_AOJI01000037.1 | contig |
| *Halorubrum sodomense* | NZ_FOYN01000001.1 - NZ_FOYN01000009.1 | scaffold |
| *Halorubrum salinarum* | NZ_CP053941.1, NZ_CP053942.1, NZ_CP053943.1 | chromosome, two plasmids pHAR01, pHAR02 |
| *Halorubrum tropicale* | NZ_LIST01000001.1 - NZ_LIST01000017.1 | contig |
| *Halorubrum terreste* | NZ_AOIW01000001.1 - NZ_AOIW01000079.1 | contig |
| *Halorubrum distributum* | NZ_AOJN01000001.1 - NZ_AOJN01000068.1 | contig |
| *Halorubrum arcis* | NZ_AOJJ01000001.1 - NZ_AOJJ01000110.1 | contig |
| *Halorubrum litoreum* | NZ_AOJF01000001.1 - NZ_AOJF01000063.1 | contig |
| *Halorubrum ruber* | NZ_CP073695.1 | chromosome |
| *Halorubrum xinjiangense* | NZ_FNBO01000001.1 - NZ_FNBO01000025.1 | scaffold |
| *Halorubrum californiense* | NZ_AOJK01000001.1 - NZ_AOJK01000083.1 | contig |
| *Halorubrum trapanicum* | NZ_JAGGKE010000001.1 - NZ_JAGGKE010000030.1 | contig |
| *Halorubrum coriense* | NZ_AOJL01000001.1 - NZ_AOJL01000069.1 | contig |

| | | |
|---|---|---|
| *Halorubrum ezzemoulense* | NZ_CP034940.1, NZ_CP034941.1, NZ_CP034942.1 | Chromosome, megaplasmid, plasmid |
| *Halorubrum hochsteinianum* | NZ_CP098415.1, NZ_CP098416.1 | Chromosome, plasmid |
| *Halorubrum tebenquichense* | NZ_AOJD01000001.1 - NZ_AOJD01000093.1 | contig |
| *Halorubrum ejinorense* | NZ_BAAADQ010000001.1 - NZ_BAAADQ010000021.1 | scaffold |
| *Halorubrum alkaliphilum* | NZ_JAGGKQ010000001.1 - NZ_JAGGKQ010000064.1 | contig |
| *Halorubrum aquaticum* | NZ_FOPZ01000001.1 - NZ_FOPZ01000040.1 | contig |
| *Halorubrum rubrum* | NZ_JANHDM010000001.1 - NZ_JANHDM010000025.1 | scaffold |
| *Halorubrum halodurans* | NZ_NHPJ01000001.1 - NZ_NHPJ01000143.1 | contig |
| *Halorubrum cibi* | NZ_FXTD01000001.1 - NZ_FXTD01000035.1 | contig |
| *Halorubrum vacuolatum* | NZ_FZNQ01000001.1 - NZ_FZNQ01000072.1 | contig |
| *Halorubrum aethiopicum* | NZ_LOAJ01000001.1 - NZ_LOAJ01000003.1 | contig |
| *Haloparvum sedimenti* | NZ_LKIR01000001.1 - NZ_LKIR01000143.1 | scaffold |
| *Halopenitus malekzadehii* | NZ_FNWU01000001.1 - NZ_FNWU01000041.1 | scaffold |
| *Halopenitus persicus* | NZ_AP017558.1, NZ_AP017559.1 | chromosome, plasmid pCBA1233 |
| *Halolamina litorea* | NZ_JANHGR010000001.1 - NZ_JANHGR010000004.1 | scaffold |
| *Halolamina pelagica* | NZ_FOXI01000001.1 - NZ_FOXI01000037.1 | contig |
| *Halolamina rubra* | NZ_BBJN01000001.1 - NZ_BBJN01000071.1 | contig |
| *Halolamina sediminis* | NZ_CVUA01000001.1 | contig |
| *Halolamina salifodinae* | NZ_JAGGLC010000001.1 - NZ_JAGGLC010000009.1 | contig |
| *Halonotius terrestis* | NZ_RKLU01000001.1 - NZ_RKLU01000026.1 | scaffold |
| *Halonotius roseus* | NZ_SESI01000001.1 - NZ_SESI01000004.1 | scaffold |
| *Halalkaliarchaeum desulfuricum* | NZ_CP025066.1 | chromosome |
| *Halohasta litorea* | NZ_JANHDJ010000001.1 - NZ_JANHDJ010000034.1 | scaffold |
| *Halohasta litchfieldiae* | NZ_CP024845.1 | chromosome |
| *Halohasta salina* | NZ_JAMZIE010000001.1 - NZ_JAMZIE010000025.1 | scaffold |
| *Halonotius pteroides* | NZ_QMDW01000001.1 - NZ_QMDW01000115.1 | contig |
| *Halonotius aquaticus* | NZ_QKNY01000001.1 - NZ_QKNY01000018.1 | scaffold |
| *Natronocalculus amylovorans* | NZ_JAKRVX010000001.1 - NZ_JAKRVX010000020.1 | contig |
| *Halalkalirubrum salinum* | NZ_SRSG01000001.1 - NZ_SRSG01000062.1 | scaffold |
| *Halopelagius inordinatus* | NZ_FOOQ01000001.1 - NZ_FOOQ01000014.1 | contig |
| *Halopelagius longus* | NZ_FNKQ01000001.1 - NZ_FNKQ01000009.1 | contig |
| *Halobellus rarus* | NZ_JANHDI010000001.1 - NZ_JANHDI010000025.1 | scaffold |

| | | |
|---|---|---|
| *Halobellus litoreus* | NZ_JANHAW010000001.1 - NZ_JANHAW010000010.1 | scaffold |
| *Halobellus limi* | NZ_FNVN01000001.1 - NZ_FNVN01000011.1 | scaffold |
| *Halobellus rufus* | NZ_BBJO01000001.1 - NZ_BBJO01000081.1 | contig |
| *Halobellus captivus* | NZ_VJXQ01000001.1 - NZ_VJXQ01000021.1 | scaffold |
| *Halobellus inordinatus* | NZ_CP101825.1, NZ_CP101826.1 | chromosome, plasmid |
| *Halobellus clavatus* | NZ_FNPB01000001.1 - NZ_FNPB01000030.1 | contig |
| *Halobellus ruber* | NZ_JACKXD010000001.1 - NZ_JACKXD010000012.1 | contig |
| *Halobellus salinus* | NZ_BMOC01000001.1 - NZ_BMOC01000054.1 | scaffold |
| *Halogeometricum borinquense* | NC_014729.1, NC_014731.1, NC_014732.1, NC_014735.1, NC_014736.1, NC_014737.1 | chromosome, plasmids pHBOR02, pHBOR04, pHBOR1, pHBOR03, pHBOR05 |
| *Halogeometricum pallidum* | NZ_AOIV01000001.1 - NZ_AOIV01000045.1 | contig |
| *Halogeometricum limi* | NZ_FOYS01000001.1 - NZ_FOYS01000008.1 | scaffold |
| *Halogeometricum rufum* | NZ_FOYT01000001.1 - NZ_FOYT01000007.1 | contig |
| *Haloquadratum walsbyi* | NC_088212.1 NC_008213.1 | chromosome, plasmid PL47 |
| *Haloferax larsenii* | NZ_CP078063.1, NZ_CP078064.1, NZ_CP078065.1 | chromosome, two plasmids pHl5678-1, pHl5678-2 |
| *Haloferax elongans* | NZ_AOLK01000001.1 - NZ_AOLK01000027.1 | contig |
| *Haloferax litoreum* | NZ_WKJO01000001.1 - NZ_WKJO01000003.1 | contig |
| *Haloferax profundi* | NZ_LOPV01000001.1 - NZ_LOPV01000716.1 | contig |
| *Haloferax marinisediminis* | NZ_WKJP01000001.1 - NZ_WKJP01000006.1 | contig |
| *Haloferax marinum* | NZ_WKJQ01000001.1- NZ_WKJQ01000003.1 | contig |
| *Haoferax massiliensis* | NZ_CSTE01000001.1 - NZ_CSTE01000008.1 | scaffold |
| *Haloferax sulfurifontis* | NZ_BMCI01000001.1 - NZ_BMCI01000021.1 | contig |
| *Haloferax denitrificans* | NZ_AOLP01000001.1 - NZ_AOLP01000021.1 | contig |
| *Haloferax marisrubi* | NZ_LOPW02000001.1 - NZ_LOPW02000026.1 | scaffold |
| *Haloferax gibbonsii* | NZ_CP011947.1 - NZ_CP011951.1 | chromosome, plasmids pHG1, pHG2, pHG3 and pHG4 |
| *Haloferax prahovense* | NZ_LK053000.1 - NZ_LK053004.1 | scaffold |
| *Haloferax mediterranei* | NZ_CP039139.1 - NZ_CP039142.1 | chromosome, plasmids pHME505, pHME322 and pHME132 |
| *Haloferax mucosum* | NZ_AOLN01000001.1 - NZ_AOLN01000020.1 | conting |
| *Haloferax volcanii* DS2 | NC_013964.1- NC_013968.1 | chromosome, four plasmids pHV1, pHV2, pHV3, pHV4 |
| *Salinigranum rubrum* | NZ_CP026309.1, NZ_CP026310.1 - NZ_CP026314.1 | chromosome, five plasmids |

| | | |
|---|---|---|
| *Salinigranum halophilum* | NZ_SSNL01000003.1 - NZ_SSNL01000012.1 | scaffold |
| *Salinigranum marinum* | NZ_CP100461.1, NZ_CP100462.1 - NZ_CP100464.1 | chromosome, three plasmids |
| *Salinigranum salinum* | NZ_VTOJ01000001.1 - NZ_VTOJ01000005.1 | scaffold |
| *Haloplanus vescus* | NZ_FNQT01000001.1 - NZ_FNQT01000010.1 | scaffold |
| *Haloplanus rallus* | NZ_CP034344.1, NZ_CP034345.1 | chromosome, plasmid |
| *Haloplanus natans* | NZ_KE386573.1, NZ_KE386574.1 | scaffold |
| *Haloplanus salinarum* | NZ_CP101823.1 | chromosome |
| *Haloplanus ruber* | NZ_JANHDK010000001.1 - NZ_JANHDK010000009.1 | scaffold |
| *Haloplanus aerogenes* | NZ_CP034145.1, NZ_CP034146.1 | chromosome, plasmid pJCM16430-01 |
| *Haloplanus salinus* | NZ_QPHM01000001.1 - NZ_QPHM01000004.1 | contig |
| *Haloplanus rubicundus* | NZ_CP031147.1, NZ_CP031148.1, NZ_CP031149.1 | chromosome, two plasmids pCBA1112-01, pCBA1112-02 |
| *Salinirubrum litoreum* | NZ_JAJCVJ010000001.1 - NZ_JAJCVJ010000005.1 | Scaffold |

**Table S3. Properties of HGTV-1 encoded tRNAs based on tRNAScan-SE 2.0 prediction.**

| tRNA | AA | anticodon | codon | CG% | Length (nt) | location |
|---|---|---|---|---|---|---|
| Hgr_chr.trna1 | Ala | GGC | GCC | 62,5 | 72 | 180042-180113(+) |
| Hgr_chr.trna2 | Ala | GGC | GCC | 62,5 | 72 | 182277-182348(+) |
| Hgr_chr.trna3 | Thr | CGT | ACG | 58,9 | 73 | 311066-311138(+) |
| Hgr_chr.trna4 | iMet | CAT | ATG | 58,7 | 75 | 451475-451549(+) |
| Hgr_chr.trna5 | Tyr | GTA | TAC | 63,5 | 74 | 731201-731274(+) |
| Hgr_chr.trna6 | Asn | GTT | AAC | 64,4 | 73 | 779960-780032(+) |
| Hgr_chr.trna7 | Asn | GTT | AAC | 63 | 73 | 780245-780317(+) |
| Hgr_chr.trna8 | Ile2 | CAT | ATG | 57,3 | 75 | 780421-780495(+) |
| Hgr_chr.trna9 | Phe | GAA | TTC | 62,2 | 74 | 783574-783647(+) |
| Hgr_chr.trna10 | Gln | CTG | CAG | 56,2 | 73 | 1264806-1264878(+) |
| Hgr_chr.trna11 | Ala | TGC | GCA | 61,1 | 72 | 1347112-1347183(+) |
| Hgr_chr.trna12 | Cys | GCA | TGC | 67,1 | 76 | 1350940-1351015(+) |
| Hgr_chr.trna13 | Lys | TTT | AAA | 65,2 | 115 | 1408966-1409080(+) |
| Hgr_chr.trna14 | Gly | CCC | GGG | 64,8 | 71 | 1489885-1489955(+) |
| Hgr_chr.trna15 | Pro | CGG | CCG | 63 | 73 | 2144208-2144280(+) |
| Hgr_chr.trna16 | Leu | TAA | TTA | 61,2 | 85 | 2236881-2236965(+) |
| Hgr_chr.trna17 | Met | CAT | ATG | 68,2 | 179 | 2496342-2496520(+) |
| Hgr_chr.trna18 | Pro | GGG | CCC | 62 | 71 | 2549043-2549113(+) |
| Hgr_chr.trna19 | Val | CAC | GTG | 64 | 75 | 2577970-2578044(+) |
| Hgr_chr.trna20 | Ser | GCT | AGC | 67,1 | 85 | 2680276-2680360(+) |
| Hgr_chr.trna21 | Leu | CAG | CTG | 61,2 | 85 | 2682724-2682808(+) |
| Hgr_chr.trna22 | Thr | TGT | ACA | 56,8 | 74 | 2784208-2784281(+) |
| Hgr_chr.trna23 | Glu | TTC | GAA | 58,7 | 75 | 2794703-2794777(+) |
| Hgr_chr.trna24 | Leu | CAA | TTG | 58,3 | 84 | 2808471-2808554(+) |
| Hgr_chr.trna25 | Glu | CTC | GAG | 56 | 75 | 2898405-2898479(+) |
| Hgr_chr.trna26 | Tyr | GTA | TAC | 62,2 | 74 | 2949002-2949075(+) |

| | | | | | | |
|---|---|---|---|---|---|---|
| Hgr_chr.trna27 | Leu | TAG | CTA | 64,3 | 84 | 3046128-3046211(+) |
| Hgr_chr.trna28 | Arg | TCG | CGA | 57,3 | 75 | 3348716-3348790(+) |
| Hgr_chr.trna29 | Trp | CCA | TGG | 61,1 | 175 | 3608042-3608216(-) |
| Hgr_chr.trna30 | Lys | CTT | AAG | 62,2 | 74 | 3007549-3007622(-) |
| Hgr_chr.trna31 | Ala | TGC | GCA | 61,1 | 72 | 2929012-2929083(-) |
| Hgr_chr.trna32 | Ser | TGA | TCA | 63,4 | 93 | 2785141-2785233(-) |
| Hgr_chr.trna33 | Ser | TGA | TCA | 59 | 83 | 2760414-2760496(-) |
| Hgr_chr.trna34 | Leu | GAG | CTC | 67,1 | 85 | 2756530-2756614(-) |
| Hgr_chr.trna35 | Ile | GAT | ATC | 62,2 | 74 | 2449948-2450021(-) |
| Hgr_chr.trna36 | Val | GAC | GTC | 56 | 75 | 2341782-2341856(-) |
| Hgr_chr.trna37 | Val | GAC | GTC | 56 | 75 | 2341666-2341740(-) |
| Hgr_chr.trna38 | Gly | GCC | GGC | 50,7 | 71 | 2301983-2302053(-) |
| Hgr_chr.trna39 | Gly | GCC | GGC | 50,7 | 71 | 2301894-2301964(-) |
| Hgr_chr.trna40 | Thr | GGT | ACC | 61,1 | 72 | 1487255-1487326(-) |
| Hgr_chr.trna41 | Gln | TTG | CAA | 60,7 | 89 | 1434202-1434290(-) |
| Hgr_chr.trna42 | Arg | CCG | CGG | 63 | 73 | 1355316-1355388(-) |
| Hgr_chr.trna43 | Arg | TCT | AGA | 57,3 | 75 | 1115952-1116026(-) |
| Hgr_chr.trna44 | His | GTG | CAC | 61,6 | 73 | 1053015-1053087(-) |
| Hgr_chr.trna45 | His | GTG | CAC | 60 | 65 | 1036648-1036712(-) |
| Hgr_chr.trna46 | Pro | TGG | CCA | 65,8 | 73 | 1027725-1027797(-) |
| Hgr_chr.trna47 | Ser | GGA | TCC | 60,5 | 81 | 903983-904063(-) |
| Hgr_chr.trna48 | Ala | CGC | GCG | 61,1 | 72 | 851341-851412(-) |
| Hgr_chr.trna49 | Asp | GTC | GAC | 64,4 | 73 | 801487-801559(-) |
| Hgr_chr.trna50 | Asp | GTC | GAC | 64,4 | 73 | 800618-800690(-) |
| Hgr_chr.trna51 | Asp | GTC | GAC | 64,4 | 73 | 800500-800572(-) |
| Hgr_chr.trna52 | Arg | GCG | CGC | 58,9 | 73 | 774306-774378(-) |
| Hgr_chr.trna53 | Val | TAC | GTA | 60,8 | 74 | 766470-766543(-) |
| Hgr_chr.trna54 | Ser | CGA | TCG | 54,7 | 86 | 616533-616618(-) |
| Hgr_chr.trna55 | Gln | TTG | CAA | 60,3 | 73 | 336496-336568(-) |
| Hgr_chr.trna56 | Gly | TCC | GGA | 59,2 | 71 | 198773-198843(-) |
| Hgr_chr.trna57 | Arg | CCT | AGG | 65,8 | 73 | 193123-193195(-) |
| Hgr_p222.trna1 | iMet | CAT | ATG | 56 | 75 | 7711-7785(+) |
| Hgr_p222.trna2 | Ala | TGC | GCA | 61,1 | 72 | 82366-82437(-) |
| HGTV.trna1 | Tyr | GTA | TAC | 38,6 | 57 | 75772-75828(-) |
| HGTV.trna2 | Undet | NNN | NNN | 52,3 | 132 | 70440-70571(-) |
| HGTV.trna3 | Arg | GCG | CGC | 51,4 | 72 | 53349-53420(-) |
| HGTV.trna4 | Asp | GTC | GAC | 59,2 | 71 | 52305-52375(-) |
| HGTV.trna5 | Glu | TTC | GAA | 55,7 | 88 | 52081-52168(-) |
| HGTV.trna6 | Phe | GAA | TTC | 52,7 | 74 | 51637-51710(-) |
| HGTV.trna7 | Glu | CTC | GAG | 57,9 | 76 | 51557-51632(-) |
| HGTV.trna8 | Gly | TCC | GGA | 60,6 | 71 | 50549-50619(-) |
| HGTV.trna9 | Gly | GCC | GGC | 62 | 71 | 50474-50544(-) |
| HGTV.trna10 | Asn | GTT | AAC | 54,8 | 73 | 50351-50423(-) |
| HGTV.trna11 | Tyr | GTA | TAC | 56,2 | 73 | 50275-50347(-) |
| HGTV.trna12 | Lys | CTT | AAG | 52,6 | 76 | 50196-50271(-) |
| HGTV.trna13 | Met | CAT | ATG | 55,8 | 86 | 50108-50193(-) |
| HGTV.trna14 | Ala | TGC | GCA | 54,4 | 103 | 49226-49328(-) |
| HGTV.trna15 | Thr | TGT | ACA | 61,4 | 57 | 49039-49095(-) |
| HGTV.trna16 | Thr | CGT | ACG | 59,5 | 74 | 48955-49028(-) |
| HGTV.trna17 | Thr | GGT | ACC | 53,5 | 71 | 48881-48951(-) |

| | | | | | | |
|---|---|---|---|---|---|---|
| HGTV.trna18 | Lys | TTT | AAA | 61,6 | 73 | 48804-48876(-) |
| HGTV.trna19 | Gln | TTG | CAA | 63 | 73 | 48548-48620(-) |
| HGTV.trna20 | Trp | CCA | TGG | 52,1 | 71 | 48474-48544(-) |
| HGTV.trna21 | Ser | CGA | TCG | 58,8 | 85 | 48384-48468(-) |
| HGTV.trna22 | Undet | NNN | NNN | 54,2 | 83 | 48164-48246(-) |
| HGTV.trna23 | Gln | CTG | CAG | 53,4 | 73 | 48089-48161(-) |
| HGTV.trna24 | Ile | GAT | ATC | 49,3 | 75 | 48011-48085(-) |
| HGTV.trna25 | Cys | GCA | TGC | 63,9 | 72 | 47538-47609(-) |
| HGTV.trna26 | Arg | TCG | CGA | 56,8 | 74 | 46701-46774(-) |
| HGTV.trna27 | Arg | TCT | AGA | 53,4 | 73 | 46357-46429(-) |
| HGTV.trna28 | Leu | TAG | CTA | 53,6 | 84 | 45786-45869(-) |
| HGTV.trna29 | Leu | GAG | CTC | 57,1 | 84 | 45617-45700(-) |
| HGTV.trna30 | Leu | CAA | TTG | 54,2 | 83 | 45531-45613(-) |
| HGTV.trna31 | Leu | TAA | TTA | 55,3 | 85 | 45442-45526(-) |
| HGTV.trna32 | Pro | TGG | CCA | 57,7 | 71 | 42770-42840(-) |
| HGTV.trna33 | Val | TAC | GTA | 44 | 75 | 42688-42762(-) |
| HGTV.trna34 | Val | GAC | GTC | 50,7 | 75 | 42609-42683(-) |

**Table S4. Codon usage in *Halogranum roseipondis* SS5-1$^{T}$ and HGTV-1 genomes**

| Codon | Amino acid | *Halogranum roseipondis* Fraction | Frequency | Number | HGTV-1 Fraction | Frequency | Number |
|---|---|---|---|---|---|---|---|
| GCA | Ala | 0,138 | 11,54 | 19911 | 0,256 | 14,994 | 719 |
| GCC | Ala | 0,28 | 23,326 | 40247 | 0,242 | 14,16 | 679 |
| GCG | Ala | **0,404** | **33,679** | **58111** | **0,268** | **15,704** | **753** |
| GCT | Ala | 0,179 | 14,905 | 25718 | 0,233 | 13,639 | 654 |
| TGC | Cys | **0,521** | **11,022** | **19018** | **0,549** | **12,638** | **606** |
| TGT | Cys | 0,479 | 10,116 | 17454 | 0,451 | 10,386 | 498 |
| GAC | Asp | **0,724** | **33,79** | **58301** | **0,574** | **23,086** | **1107** |
| GAT | Asp | 0,276 | 12,852 | 22175 | 0,426 | 17,142 | 822 |
| GAA | Glu | 0,391 | 16,069 | 27726 | **0,53** | **26,047** | **1249** |
| GAG | Glu | **0,609** | **25,019** | **43169** | 0,47 | 23,065 | 1106 |
| TTC | Phe | **0,781** | **15,866** | **27376** | **0,672** | **25,505** | **1223** |
| TTT | Phe | 0,219 | 4,439 | 7659 | 0,328 | 12,45 | 597 |
| GGA | Gly | 0,24 | 18,227 | 31450 | **0,308** | **17,914** | **859** |
| GGC | Gly | **0,306** | **23,26** | **40134** | 0,25 | 14,515 | 696 |
| GGG | Gly | 0,213 | 16,215 | 27977 | 0,165 | 9,572 | 459 |
| GGT | Gly | 0,24 | 18,249 | 31487 | 0,277 | 16,121 | 773 |
| CAC | His | **0,668** | **16,129** | **27829** | **0,525** | **14,848** | **712** |
| CAT | His | 0,332 | 8,034 | 13862 | 0,475 | 13,41 | 643 |
| ATA | Ile | 0,176 | 3,645 | 6290 | 0,284 | 13,764 | 660 |
| ATC | Ile | **0,632** | **13,117** | **22633** | 0,342 | 16,579 | 795 |
| ATT | Ile | 0,193 | 3,998 | 6899 | **0,374** | **18,144** | **870** |
| AAA | Lys | 0,346 | 4,357 | 7517 | 0,366 | 11,512 | 552 |
| AAG | Lys | **0,654** | **8,222** | **14187** | **0,634** | **19,958** | **957** |
| CTA | Leu | 0,077 | 4,725 | 8153 | 0,096 | 8,363 | 401 |
| CTC | Leu | **0,413** | **25,489** | **43979** | **0,262** | **22,732** | **1090** |
| CTG | Leu | 0,229 | 14,161 | 24434 | 0,136 | 11,762 | 564 |
| CTT | Leu | 0,129 | 7,959 | 13732 | 0,213 | 18,519 | 888 |
| TTA | Leu | 0,037 | 2,277 | 3929 | 0,119 | 10,344 | 496 |
| TTG | Leu | 0,116 | 7,139 | 12318 | 0,174 | 15,057 | 722 |
| ATG | Met | 1 | 8,204 | 14155 | 1 | 13,973 | 670 |
| AAC | Asn | **0,772** | **13,432** | **23175** | **0,508** | **17,893** | **858** |
| AAT | Asn | 0,228 | 3,963 | 6837 | 0,492 | 17,351 | 832 |
| CCA | Pro | 0,164 | 12,559 | 21670 | 0,269 | 13,66 | 655 |
| CCC | Pro | 0,215 | 16,406 | 28308 | 0,207 | 10,511 | 504 |
| CCG | Pro | **0,453** | **34,642** | **59771** | **0,272** | **13,827** | **663** |
| CCT | Pro | 0,168 | 12,831 | 22138 | 0,252 | 12,784 | 613 |

| | | | | | | | |
|---|---|---|---|---|---|---|---|
| CAA | Gln | 0,346 | 7,47 | 12888 | **0,505** | **13,806** | **662** |
| CAG | Gln | **0,654** | **14,144** | **24405** | 0,495 | 13,535 | 649 |
| AGA | Arg | 0,077 | 13,159 | 22705 | 0,15 | 14,869 | 713 |
| AGG | Arg | 0,074 | 12,57 | 21689 | 0,128 | 12,721 | 610 |
| CGA | Arg | **0,267** | **45,623** | **78719** | **0,236** | **23,441** | **1124** |
| CGC | Arg | 0,198 | 33,893 | 58480 | 0,152 | 15,057 | 722 |
| CGG | Arg | 0,202 | 34,5 | 59527 | 0,14 | 13,91 | 667 |
| CGT | Arg | 0,182 | 31,168 | 53777 | 0,193 | 19,186 | 920 |
| AGC | Ser | 0,131 | 14,715 | 25390 | 0,12 | 11,908 | 571 |
| AGT | Ser | 0,101 | 11,387 | 19647 | 0,147 | 14,619 | 701 |
| TCA | Ser | 0,1 | 11,193 | 19312 | 0,147 | 14,598 | 700 |
| TCC | Ser | 0,16 | 17,957 | 30983 | 0,169 | 16,746 | 803 |
| TCG | Ser | **0,392** | **44,048** | **76001** | **0,256** | **25,463** | **1221** |
| TCT | Ser | 0,117 | 13,143 | 22677 | 0,161 | 15,975 | 766 |
| ACA | Thr | 0,143 | 10,234 | 17658 | 0,182 | 11,637 | 558 |
| ACC | Thr | 0,258 | 18,441 | 31818 | 0,214 | 13,701 | 657 |
| ACG | Thr | **0,437** | **31,181** | **53801** | **0,34** | **21,772** | **1044** |
| ACT | Thr | 0,161 | 11,495 | 19834 | 0,263 | 16,851 | 808 |
| GTA | Val | 0,122 | 8,685 | 14986 | 0,227 | 15,495 | 743 |
| GTC | Val | **0,472** | **33,49** | **57784** | **0,308** | **20,98** | **1006** |
| GTG | Val | 0,222 | 15,723 | 27128 | 0,206 | 14,014 | 672 |
| GTT | Val | 0,184 | 13,017 | 22460 | 0,259 | 17,685 | 848 |
| TGG | Trp | 1 | 12,518 | 21599 | 1 | 13,993 | 671 |
| TAC | Tyr | **0,7** | **8,813** | **15206** | **0,552** | **17,205** | **825** |
| TAT | Tyr | 0,3 | 3,778 | 6518 | 0,448 | 13,952 | 669 |
| TAA | | 0,123 | 2,182 | 3764 | 0,309 | 10,803 | 518 |
| TAG | | 0,251 | 4,46 | 7695 | 0,215 | 7,529 | 361 |
| TGA | | 0,627 | 11,148 | 19235 | 0,476 | 16,621 | 797 |
